# TAILORED DC INDUCE PROTECTIVE HIV-1 SPECIFIC POLYFUNCTIONAL CD8+ T CELLS IN THE LYMPHOID TISSUE FROM HUMANIZED BLT MICE

**DOI:** 10.1101/2021.06.30.450486

**Authors:** Marta Calvet-Mirabent, Daniel T. Claiborne, Maud Deruaz, Serah Tanno, Carla Serra, Cristina Delgado-Arévalo, Ildefonso Sánchez-Cerrillo, Ignacio de los Santos, Jesús Sanz, Lucio García-Fraile, Francisco Sánchez-Madrid, Arantzazu Alfranca, María Ángeles Muñoz-Fernández, Todd M. Allen, Maria J. Buzón, Alejandro Balazs, Vladimir Vrbanac, Enrique Martín-Gayo

## Abstract

Effective function of CD8^+^ T cells and enhanced innate activation of dendritic cells (DC) in response to HIV-1 is linked to protective antiviral immunity in controllers. Manipulation of DC targeting the master regulator TANK-binding Kinase 1 (TBK1) might be useful to acquire controller-like properties. Here, we evaluated the impact of TBK1-primed DC inducing protective CD8^+^ T cell responses in lymphoid tissue and peripheral blood and their association with reduced HIV-1 disease progression *in vivo* in the humanized bone marrow, liver and thymus (hBLT) mouse model. A higher proportion of hBLT-mice vaccinated with TBK1-primed DC exhibited less severe CD4^+^ T cell depletion following HIV-1 infection compared to control groups. This was associated with infiltration of CD8^+^ T cells in the white pulp from the spleen, reduced spread of infected p24^+^ cells to secondary lymphoid organs and with preserved abilities of CD8^+^ T cells from the spleen and blood of vaccinated animals to induce specific polyfunctional responses upon antigen stimulation. Therefore, TBK1-primed DC might be an useful tool for subsequent vaccine studies.

**Author summary:** Emulating protective immunological characteristics from individuals capable of spontaneously controlling HIV-1 infection might be useful for the development of a protective vaccine. Enhanced function of dendritic cells (DC) in these HIV-1 controllers depends on the activation of TANK-binding Kinase 1 (TBK1) and might associate with protective T cells. Our study evaluated the ability of DCs trained through TBK1 activation inducing protective adaptive immune responses against HIV-1 and reducing disease progression *in vivo*, using a humanized mouse model. Our data indicate that mice vaccinated with tailored DC exhibit delayed disease progression, increased induction of protective CD8+ T lymphocyte subsets in the lymphoid tissue and blood upon antigen recognition. Therefore, trained-DC might be an useful tool for future HIV-1 vaccine designs.

## Introduction

A remaining challenge to end the HIV-1 pandemic is the development of an effective vaccine capable of providing protective and long-lasting immunity against HIV-1 infection. While previous efforts to achieve this goal have failed (1, 2), the scientific community has come to understand that the induction of effective and durable HIV-1-specific T cell responses in different anatomical compartments will most likely require the targeting and fine-tuning of specific innate immune cell subsets, such as dendritic cells (DC). DC play a critical role during the priming of specific adaptive immune responses, since they are capable of both efficiently presenting antigens (Ags) to T cells and also mediating the polarization of effector lymphocytes (3-7). In fact, DC-based therapeutic vaccines have shown very promising results in clinical trials for cancer therapy (8). However, although encouraging, previous DC-based HIV-1 vaccination strategies have demonstrated limited abilities priming durable memory HIV-1-specific T cell responses (9-13). In addition, most vaccine studies used adjuvants systemically as a means to globally increase innate immune activation, without considering their individual impact on specific DC functional characteristics (14).

Previous studies showed that conventional DC (cDC) from HIV-1 elite controllers (EC) are capable of efficiently detecting HIV-1 reverse transcripts (15, 16) and inducing activation of the signal transducer TANK-binding Kinase 1 (TBK1)(17, 18). This mechanism leads to enhanced capabilities to prime polyfunctional HIV-1-specific CD8^+^ T cell responses, which are associated with effective control of HIV-1 infection (19-22). Therefore, TBK1 may, in principle, represent a therapeutic target to improve DC maturation towards an EC-like phenotype and to more efficiently activate protective antiviral CD8^+^ T cell responses in a broader population of individuals. Combined stimulation of DC with ligands to multiple intracellular sensors upstream TBK1 such as cGAS, RIG-I, MDA5 or TLR3(23), could synergistically act as TBK1 adjuvants and further improve the function of these cells. Supporting this possibility, initial studies suggested that the maturation of DC in the presence of the TLR3/RIG-I ligand Poly I:C boosts HIV-1-specific T cell responses from HIV-1-infected individuals *in vitro*(24). Multiple vaccine studies have mainly focused on analyzing activation patterns on circulating HIV-1-specific T cells, despite growing evidence of the critical role of lymphoid tissue-resident T cells controlling HIV-1 or simian immunodeficiency virus (SIV)(25, 26). Therefore, it is critical to determine the efficiency and relevance of potential novel DC-vaccine strategies inducing HIV-1-specific adaptive immune responses *in vivo* in different tissue locations.

The non-human primate model has been traditionally recognized as the gold standard *in vivo* model to test HIV-1 vaccine candidates (27). However, in addition to intrinsic differences with the human organism, this *in vivo* model might not always be accessible for initial phases of vaccine candidate evaluation. Immunodeficient NOD/SCID IL2Ry^-/-^ (NSG) mice transplanted with human fetal hematopoietic stem cells, liver and thymus (here after referred to as hBLT-mouse) represent a more accessible humanized *in vivo* system that recapitulates the development of most human myeloid and lymphoid lineages (28-31). Importantly, hBLT-mice can be infected with HIV-1 and meet some aspects of HIV-1 disease progression, such as the depletion of CD4^+^ T cell lymphocytes and the induction of specific adaptive immune responses, including cytotoxic CD8^+^ T cells (CTL) (32-35). Moreover, the hBLT model supports the induction of effector memory HIV-1-specific CD8^+^ T cells similar to those observed in previous vaccine studies (12, 36, 37). Despite some limitations, the hBLT mouse represents a very attractive model for a proof-of-concept of HIV-1 vaccine study. Recent data indicate that the immunization of hBLT mice with HIV-1 Gag protein potentiates the induction of Gag-specific T cells capable of reducing HIV-1 viremia and forcing viral escape mutations (38). However, whether the hBLT model supports the induction of protective T cell responses in different lymphoid tissue compartments that could actively contribute to viral control after vaccination has not been studied in detail. In addition, little, if any, information on the polyfunctional characteristics of CD8^+^ T cells, a critical hallmark of immune control of HIV-1 infection (39, 40), has been described in this system. Finally, the impact and potential benefit of a DC-based HIV-1 vaccine on the induction of HIV-1 specific T cells and disease progression have not been tested in the hBLT mouse model yet.

In this study, we assessed the ability of TBK1-primed DC to improve parameters of immune protection against HIV-1 in the lymphoid tissue and peripheral blood using the hBTL mouse model. Our data indicate that TBK1-primed DC potentiate the infiltration of CD8^+^ T cells in the white pulp of spleen and the retention of infected HIV-1 p24^+^ cells in these areas, preventing viral spread to secondary lymphoid organs. These histological parameters induced by TBK1 DC-vaccination correlated with preserved abilities to induce polyfunctional CD8^+^ T cell responses in the spleen upon HIV-1 Gag stimulation and with less severe depletion of CD4^+^ T cells at late time points of infection in vaccinated hBLT mice. Our study provides novel evidence of enhanced cellular immunity against HIV-1 in the lymphoid tissue induced by a tailored DC-based vaccine *in vivo*, which could be useful for the development of new vaccine strategies.

## Results

### Combination of TBK1 adjuvants improves maturation and functional properties of DC

We first evaluated the efficacy of potential TBK1 adjuvants such as the TLR3 and STING agonists Poly I:C and 2′3′-di-AM(PS) to enhance maturation and functional properties of DCs. To this end, we stimulated Monocyte derived-DC (MDDC) and primary circulating CD1c^+^ cDCs with these molecules individually or in combination and monitored the phosphorylation of TBK1 and the downstream effector IRF3 as a readout of activation. As shown in Figure 1A and Supplemental Figure 1A, stimulation of both MDDC and cDC with a combination of the STING agonist and Poly:IC led to a more significant increase in TBK1 and IRF3 phosphorylation compared to individual treatments. Therefore, simultaneous stimulation with the STING agonist and Poly I:C could have significant impact on the activation and subsequent maturation of DC. To test this, we assessed expression of maturation markers and the transcription of inflammatory cytokines on primary cDCs stimulated with TBK1 adjuvants. As shown in Supplemental Figure 1B, both STING agonist and Poly I:C were able to significantly increase expression of CD40 and CD86 individually, and the combination of both TBK1 adjuvants led to limited but significant additional increase in the expression of CD40. We observed that the combination of the STING agonist and Poly I:C induced significantly higher mRNA levels of IFNβ, IL-12 and, to some extent, TNFα, suggesting an enhancement in the maturation program of cDC (Figure 1B). To determine whether these changes in cDC could be translated into improved functional antigen presenting cell properties, we first performed co-cultures of total T cells with allogeneic cDC pre-incubated in media or in the presence of different combinations of TBK1 adjuvants. cDC treated with both Poly I:C and the STING agonist were capable of inducing higher proportions of CD8^+^T cells co-expressing IFNγ and the degranulation marker CD107a (Figure 1C). Importantly, we observed that treatment of PBMC from healthy donors with Poly I:C and STING agonist in the presence of a pool of HIV-1 Gag peptides and a subsequent boost with autologous Gag-peptide loaded cDC stimulated with both TBK1 adjuvants enhanced *de novo* induction of IFNγ^+^ HIV-1 Gag-specific CD8^+^ T cells *in vitro* (Figure 1D, left). Moreover, significantly higher proportions of IFNγ^+^ CD8^+^ T co-expressing CD107a were detected in the presence of Gag-peptide loaded cDC stimulated with TBK1 adjuvants (Figure 1D, right), suggesting enhanced polyfunctionality in HIV-1-specific cells. Importantly, these effects were dependent on the presence of the Ag, since no significant increase of T cell responses was observed after stimulation only with TBK1 adjuvants (Supplemental Figure 1C). Together, these data suggest that combination of of Poly I:C and STING agonists as effective TBK1 adjuvants potentiating the maturation and function of cDCs *in vitro*.

**Figure 1.**
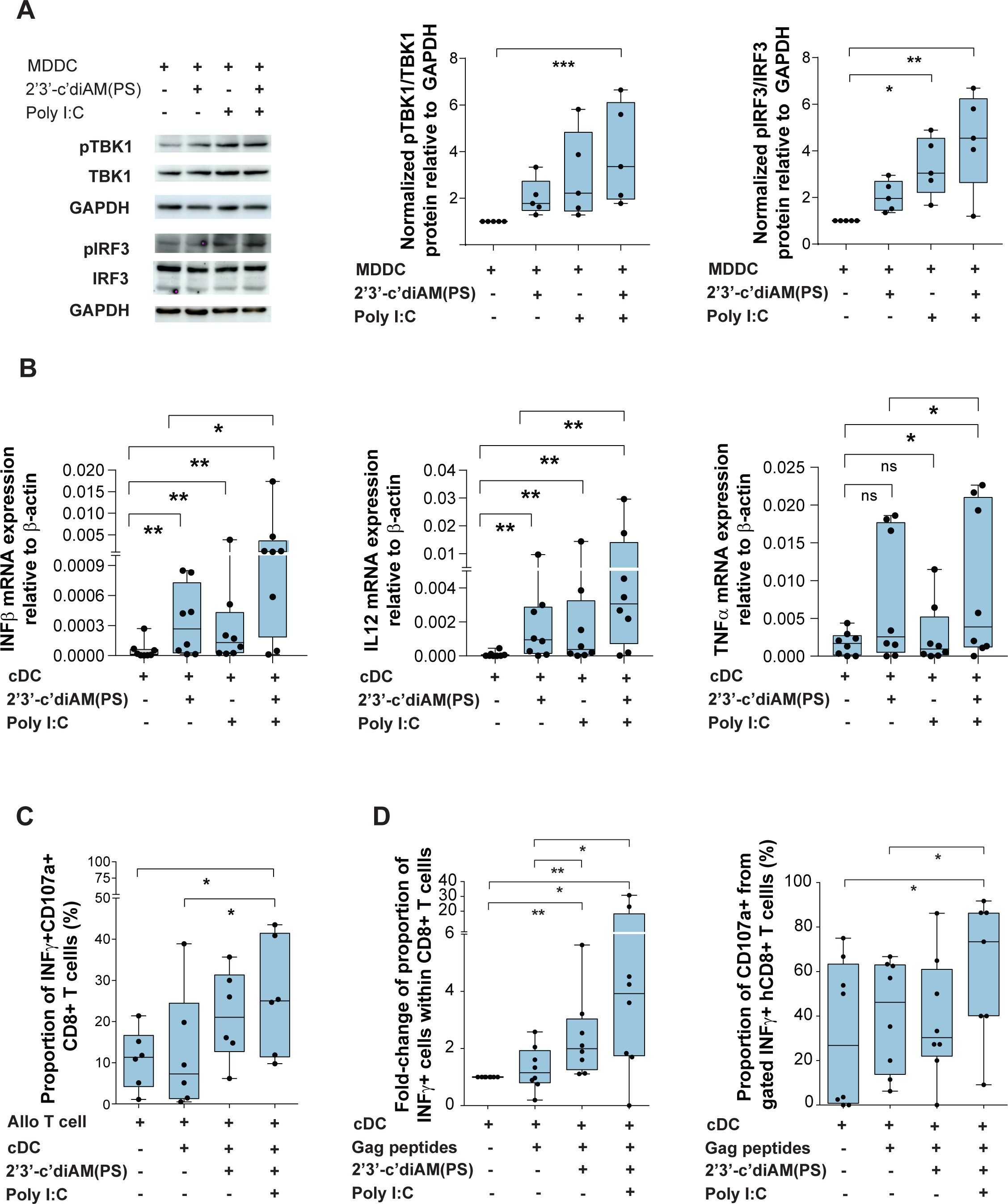
Impact of combined TBK1 adjuvants on maturation and function of DC *in vitro*. (A): Representative Western blot images of analysis of phosphorylated and total TBK1 and IRF3 proteins in Monocyte-Derived DC (MDDC) cultured in the absence or the presence of individual or combined TBK1 adjuvants (left panel). Activation of TBK1 (left plot) and IRF3 (right plot) proteins was determined by calculating the ratio of phosphorylated vs total protein and normalized to GAPDH as housekeeping protein for DCs. Data shown in the right represent ratios normalized to values from the control condition (MDDC alone) of each experiment (n=5 experiments). Statistical significance was calculated using a Kruskal-Wallis multiple comparison test with Dunn’s correction (*p<0.05; **p<0.01; ***p<0.001). (B): RT-qPCR analysis of IFNγ, IL-12 and TNFα mRNA expression normalized to β-actin levels in cDC cultured for 16 h hours with media alone or in the presence of 2’3’-c-di-AM(PS)2 and/or Poly I:C. (n=8 experiments). Statistical significance was calculated using a two-tailed matched-pairs Wilcoxon test (*p<0.05; **p<0.01). (C): Proportions of polyfunctional IFNγ^+^ CD107a^+^ CD8^+^ T cells detected by flow cytometry after culture of total T cells with allogeneic cDCs pre-treated with media or in the presence of individual or combined TBK1 adjuvants. Significance was calculated using a two-tailed Wilcoxon test (*p<0.05). (D): Proportions of *de novo* induced total (IFNγ^+^, left) and polyfunctional (IFNγ^+^ CD107a^+^, right) HIV-1-Gag-specific T cells from healthy donors cultured for 2 weeks in the absence or the presence of a pool of HIV-1 Gag peptides alone or combined with the indicated TBK1 adjuvants and restimulated with autologous cDC pre-treated in the same mentioned culture conditions. Significance was calculated using a two-tailed Wilcoxon test (*p<0.05; **p<0.01).

### Vaccination of hBLT-mice with TBK1-primed DC reduces HIV-1-mediated disease progression

We next determined whether DC activated in the presence of the TBK1 adjuvant cocktail could also induce protective responses against HIV-1 infection *in vivo* using the hBLT mouse model. To ensure that only DC were manipulated with TBK1 adjuvants, we differentiated CD11c^+^ CD14^-^ HLADR^+^ cDC and CD11c^+^ CD14^+^ HLADR^+^ MoDC-like cells *in vitro* from a portion of the human fetal CD34^+^ HSC precursors used to reconstitute the hBLT mice prior to vaccination (Supplemental Figure 2A). HSC-derived cDC and MoDC were sorted and cultured separately for 24h in the presence of media alone (MED), a pool of peptides from HIV-1 Gag alone (GAG) or in combination with our TBK1 adjuvant cocktail (GAG-ADJ) (Supplemental Figure 2B-C). The individual addition of Gag peptides did not induce significant activation of sorted cells (Supplemental Figure 2C). However, despite differences in basal expression of activation markers, both sorted DC subsets responded to the adjuvant stimulation (Supplemental Figure 2C) and cDC and MoDC from each condition were pooled for vaccination (Supplemental Figure 2D). In two experiments performed with different batches of hBLT mice, a total of n=42 hBLT animals were subdivided in 3 groups of n=14 animals that were vaccinated intravenously with either MED, GAG or GAG-ADJ DC by injection in the tail vein (Supplemental Figure 2D). Two weeks after vaccination, mice were intravenously infected with 10,000 TCID50 of JRCF HIV-1 strain. Prior pilot experiments indicated that HIV-1 plasma viremia begins to stabilize by 3 weeks post-infection (p.i.) and reaches a stable setpoint by 6 weeks p.i. in hBLT (Supplemental Figure 3A). In addition, it has been reported that at 6 weeks p.i. depletion of CD4^+^ T cells and HIV disease progression reproducibly becomes more evident in hBLT mice infected with JRCSF HIV-1(32) and is the peak time point of detection of HIV-1 specific T cell responses in the blood of these animals (38). Therefore, we analyzed clinical, histological and cellular parameters associated with protection or disease progression at three, five/six and six/seven weeks p.i. to cover these critical time points (Supplemental Figure 2D). As shown in Supplemental Figure 2E, no differences in weight were observed among the three hBLT mouse groups prior or after HIV-1 infection, suggesting vaccination did not have any significant impact on the induction of GvHD. Although all hBLT mouse groups experienced a significant reduction in circulating hCD4^+^ T cells 3 weeks after infection with HIV-1 compared to baseline (Supplemental Figure 4A, C), we observed a noticeably less severe CD4^+^ T cell depletion in the GAG-ADJ group (Figure 2A) at 5/6 weeks post infection (Figure 2A, Supplemental Figure 4A-C). Consistently, CD4^+^/CD8^+^ T cell ratio in the blood tended to be higher in GAG-ADJ mice at later time points of infection (Supplemental Figure 4B). Notably, the GAG-ADJ vaccinated group included a significantly higher proportion of animals experiencing less than 0.5-fold reduction in circulating CD4^+^ T cell numbers (CD4Hi phenotype) at these late time points (Figure 2A; Supplemental Figure 4A-C). In contrast, mice vaccinated with GAG DC experienced a dramatic depletion of CD4^+^ T cells below 0.5-fold threshold (CD4Lo phenotype) in the majority of animals from this group (Figure 2A, Supplemental Figure 4A-C). Similar results were obtained at 6/7 weeks p.i, but differences between vaccinated groups were more pronounced at 5/6 weeks p.i. (Supplemental Figure 4A-C). These effects were consistently observed in the two independent hBLT mouse batches (Supplementary Figure 4A-C). Interestingly, mice vaccinated with MED DC that had not received adjuvant or Ag were characterized by an intermediate phenotype of 50% animals exhibiting dramatic (<0.5-fold decrease) and 50% less severe (>0.5-fold decrease) depletion of CD4^+^ T cells, suggesting a partial and Ag-independent effect of vaccination with immature DC (Figure 2A, Supplemental Figure 4A-B). Interestingly, while not significant differences in plasma viremia were observed at any time point between the total 3 groups of vaccinated animals (Supplemental Figure 3B), we observed an enrichment of lower viral loads at 3 weeks p.i. in those hBLT mice displaying a less severe CD4Hi phenotype at 5/6 weeks p.i., which again were more significantly enriched in the GAG-ADJ and MED animal groups (Figure 2B). The early control of viremia seemed to be transient and no significant differences were observed in plasma viral loads by 6/7 weeks p.i (Supplemental Figure 3C). These data indicate that vaccination of hBLT mice with TBK1-trained DC is associated with less severe depletion of CD4^+^ T cells and a concomitant partial early control of HIV-1 viremia, suggesting delayed progression of HIV-1 infection in these animals.

**Figure 2.**
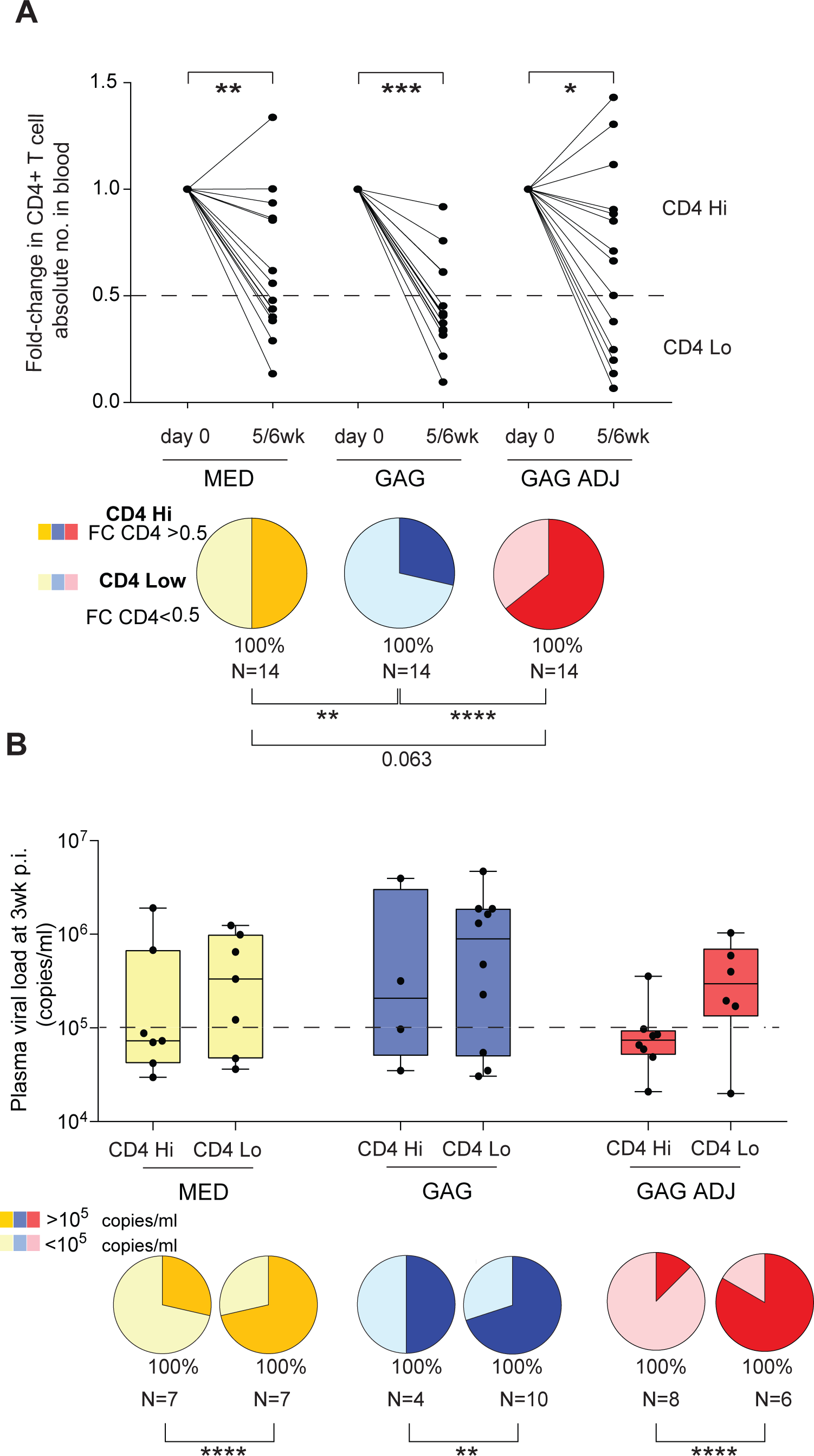
hBLT mice vaccinated with GAG-ADJ DC display less severe progression of HIV-1 infection. (A): Fold-change in circulating hCD4^+^ T cell counts in infected hBLT mice at 5-6 weeks post-infection with HIV-1 relative to basal counts present on each mouse at day 0 (upper panels). Significance was calculated using a two-tailed Wilcoxon test (*p<0.05; **p<0.01; ***p<0.001). Pie charts showing percentage of mice displaying less severe decrease of hCD4^+^ T cell counts (hCD4^+^ T cell fold change ≥ 0.5; CD4 Hi) and those animals with severe depletion (hCD4^+^ T cell fold change < 0.5; CD4 Low). Statistical significance of differences was calculated using a Chi-square test with Yates correction (**p<0.01; ****p<0.0001). (B): HIV-1 plasma viral loads (upper panels) quantified by RT-qPCR from the plasma of hBLT-mice vaccinated with MED, GAG and GAG-ADJ treated DCs at 3 weeks post infection, stratified by CD4 Hi and CD4 Low phenotypes within each indicated hBLT mouse subgroup. Pie charts (lower panels) representing mice with VL either equal or higher than 10^5^ copies/ml (dark color) or lower than 10^5^ copies/ml (light color) per treatment group and CD4^+^ T cell fold-change stratification. Statistical significance of differences was calculated using a Chi-square test with Yates correction (**p<0.01; ****p<0.0001).

**Figure 3.**
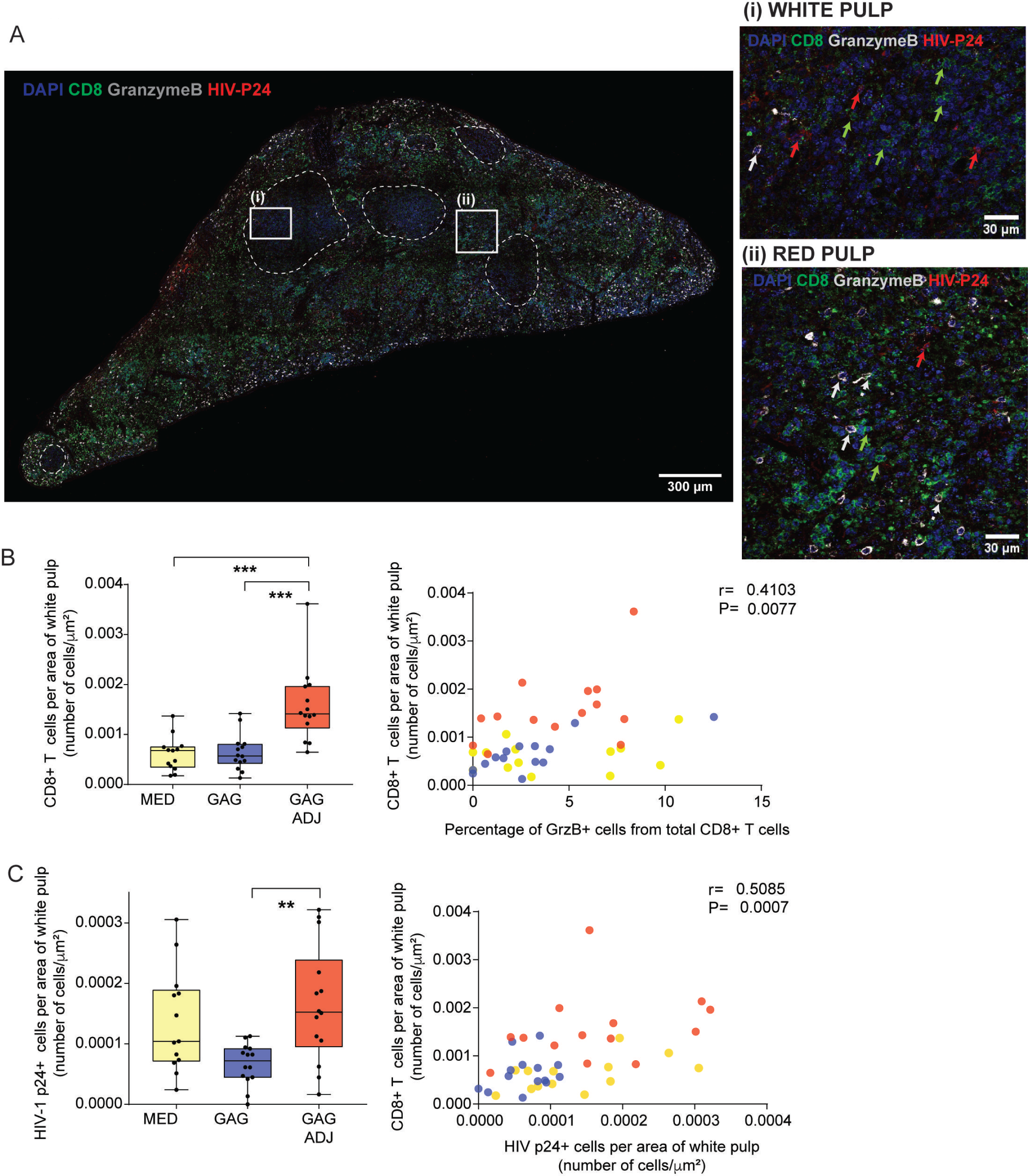
Histological analysis of CD8^+^ T cell and HIV-1-infected cell distribution in spleen from vaccinated hBLT mice. (A): Representative confocal microscopy image (magnification 40X) of a whole transversal splenic section showing staining of cell nuclei (DAPI; blue), human CD8^+^ T cells (green), Granzyme B^+^ (gray) and HIV-1 p24^+^ infected cells (red). Zoomed images (40X magnification) from selected white pulp (i) and red pulp (ii) areas highlighted by dashed lines and defined as in Supplemental Figure 4, are displayed on the right to appreciate cellular patterns. Green arrows CD8^+^ T cells; white arrow Granzyme B^+^ cell; dashed white arrow Granzyme B^+^ CD8^+^ T cells; red arrow HIV-1 p24^+^ cells. (B-C): Analysis of hCD8^+^ T cells (B, left) and HIV-1 p24^+^ cells (C, left) infiltrated in the white pulp areas from spleen of the indicated groups of hBLT mice. Significance was calculated using a Kruskal-Wallis multiple comparison test with Dunn’s correction (**p<0.01; ***p<0.001). Spearman correlation analysis of association of frequencies of CD8^+^ T cells in the white pulp and proportions of Granzyme B^+^ CD8^+^ T cells (B, right) and p24^+^ in the white pulp (C, right) are also shown. Spearman R and P values are highlighted on the upper right areas of each plot.

**Figure 4.**
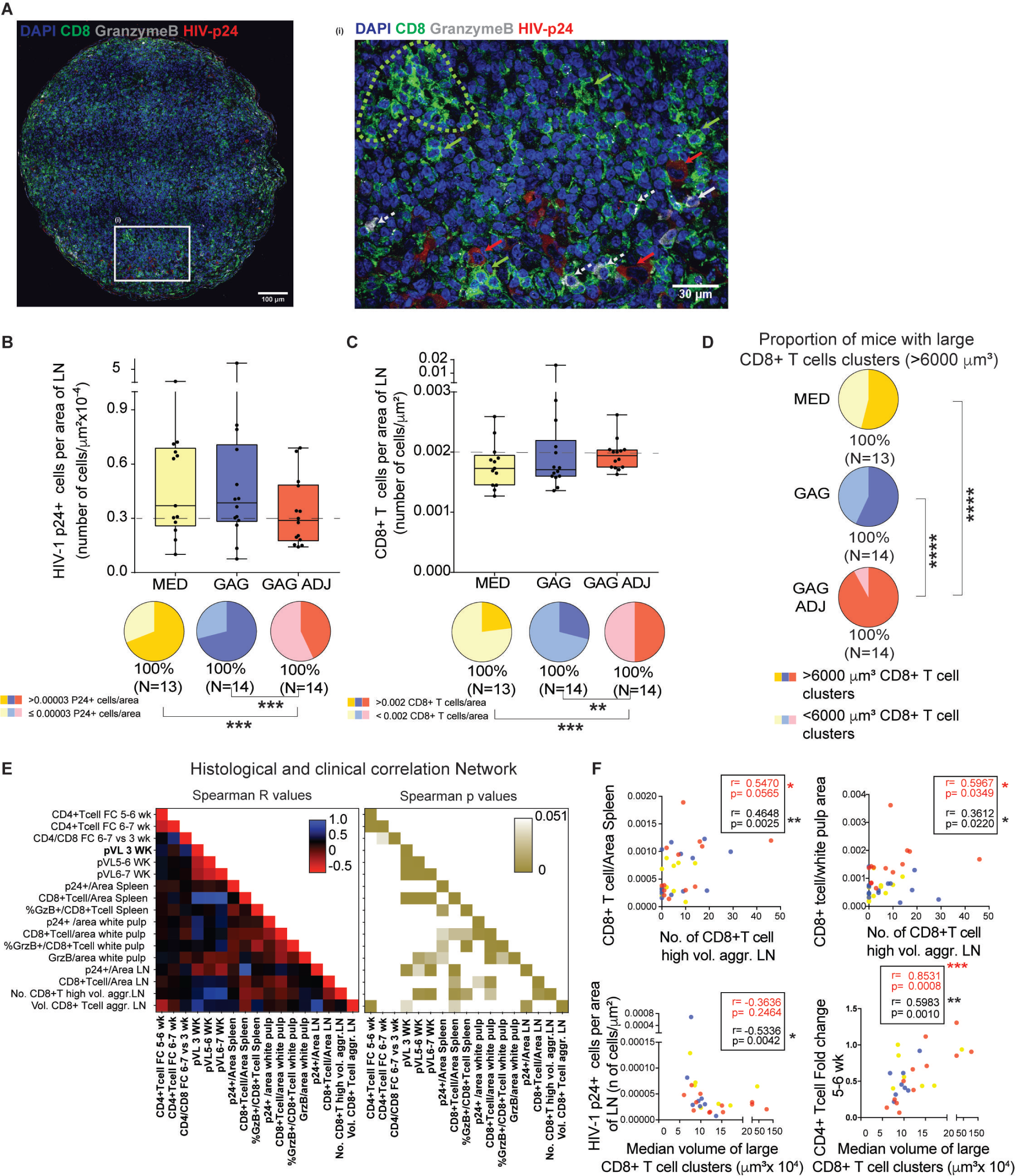
Histological CD8^+^ T cell and HIV-1 p24^+^ cell characterization in lymph nodes from vaccinated hBLT mice. (A): Representative confocal microscopy image (40x magnification) example of whole lymph node section showing staining of nuclei (DAPI, blue), hCD8^+^ T cells (green), Granzyme B^+^ (grey) and infected HIV-1 p24^+^ cells (red). A zoom of an (i) area from the same original image is shown on the right. Dashed lines highlight CD8^+^ T cell cluster areas. Green arrows CD8^+^ T cells; white arrow Granzyme B^+^ cell; dashed white arrow Granzyme B^+^ CD8^+^ T cells; red arrow HIV-1 p24^+^ cells. (B): Quantification of number of infected HIV-1 p24^+^ cells per lymph node area from the indicated hBLT mice groups. Pie charts shown below represent the percentage of mice displaying high density of infected cells per area (≥ 0.00003 p24^+^ cells/square micron) or low density of infected cells per area (< 0.00003 p24^+^ cells/square micron) within each hBLT mouse subgroup. Statistical significance of differences was calculated using a Chi-square test with Yates correction (***p<0.001). (C): Number of hCD8^+^ T cells per lymph node area (upper panel) from the indicated hBLT mice groups. Pie charts showing the percentage of mice per group displaying high density of CD8^+^ T cells per lymph node area (≥ 0.002 hCD8^+^ T cells/square micron) or low density of CD8^+^ T cells per lymph node area (< 0.002 hCD8^+^ T cells/square micron) are shown below. Statistical significance of differences in proportions of mice with enrichment of CD8^+^ T cells among groups was calculated using a Chi-square test with Yates correction (**p<0.01; ***p<0.001). (D): Percentage of mice presenting CD8^+^ T cells large volume clusters (≥ 6000 cubic microns) in the lymph nodes corresponding to the quantifications shown in Supplemental Figure 5. Statistical significance of differences was calculated using a Chi-square test with Yates correction (****p<0.0001). (E): Two tailed Spearman correlation network showing R (left heatmap) and p values (right heatmap) between selected histological parameters and plasma viral loads and fold change in PB CD4^+^ T cell count and CD4^+^/CD8^+^ T cell ratios at different times post infection. (F): Individual Spearman correlations between numbers (Upper row) and median volume (lower row) of large CD8^+^ T cell clusters (≥ 6000 cubic microns) versus the indicated spleen histological patterns (upper row) and clinical parameters (lower row). Spearman R and p values for all animals (black) and GAG-ADJ group (red) are shown on each plot.

### Accumulation of CD8^+^ T cells and HIV-1 infected cells in the white pulp after vaccination with TBK1-primed DC

To better understand differences in HIV-1 disease progression in the three groups of vaccinated hBLT mice, we analyzed histological distribution of CD8^+^ T cells and infected p24^+^ cells by immunofluorescence in tissue sections from spleen and lymph nodes (LN) from the hBLT mice at 6/7 weeks p.i. As shown in Figure 3A-B and 4A-B, p24^+^ HIV-1-infected cells could be detected in the spleen and LN of all hBLT mice, consistent with previous observations (32). No differences were observed in total HIV-1 p24^+^ cell counts in the spleen of infected animals (Supplemental Figure 5B, lower panel), and a weak enrichment on total splenic Granzyme B^+^ CD8^+^ T cells was observed in tissue sections from hBLT mice vaccinated with GAG-ADJ DC (Supplemental Figure 5B, upper panel). However, infiltrated CD8^+^ T cells were significantly higher in white pulp areas defined by hematoxylin/eosin staining, from the spleens from GAG-ADJ vaccinated hBLT mice at 6/7 weeks p.i. (Figure 3B left panel, Supplemental Figure 5A). Interestingly, increased infiltration of CD8^+^ T cells in the white pulp was significantly associated with expression of Granzyme B in the spleen (Figure 3B, right). Additionally, we observed significantly increased proportions of Granzyme B^+^ cytotoxic CD8^+^ T cells in the surrounding red pulp areas in the spleen of GAG-ADJ mice, and higher frequencies of these cells correlated with infiltration of CD8^+^ T cells in the white pulp (Supplemental Figure 5C-D). Interestingly, CD8^+^ T cells infiltrated in the spleen white pulp also tended to express higher levels of Granzyme B and CXCR5 in GAG-ADJ mice (Supplemental Figure 5C,F). In contrast, opposite patterns were observed in GAG mice (Supplemental Figure 5C,F). Notably, we also observed a significantly higher accumulation of infected p24^+^ cells in white pulp areas of spleen from these GAG-ADJ hBLT mice compared to those vaccinated with GAG, which was correlated with increased infiltration of CD8^+^ T cells in this area (Figure 3C). These effects were more significantly appreciated in CD4Hi ADJ-GAG hBLT mice displaying less severe depletion of CD4^+^ T cells compared to GAG-only hBLT mice (Supplemental Figure 5E). In contrast, reduced frequencies of HIV-1 p24^+^ cells and enrichment on CD8^+^ T cells were observed in the LN of GAG-ADJ hBLT mice and more significantly in mice from this group experiencing less severe depletion of CD4^+^ T cells (Figure 4 A-C; Supplemental Figure 6B,). In addition, numbers of p24^+^ cells per area of the LN correlated with viral load detection either at early time-points (3 weeks p.i.) and late time-points (5-6 and 6-7 weeks p.i.) and were inversely associated with CD8^+^ T cell recruitment in the spleen. (Supplemental Figure 6C; Figure 4E). Moreover, CD8^+^ T cells recruited in the LN from GAG-ADJ vaccinated hBLT animals distributed in significantly larger cell clusters (Figure 4D, Supplemental Figure 6A-D). Clustered CD8^+^ T cells did not appear preferentially express Granzyme B (Figure 4A, right). Importantly, we observed that these histological CD8^+^ T cell aggregation patterns were significantly inversely associated with less detection of p24^+^ cells in the LN and positively correlated with increased recruitment of CD8^+^ T cells to the spleen and infiltration in the white pulp areas and with less severe depletion of CD4^+^ T cells at 5/6 weeks p.i. (Figure 4E-F). Our data clearly indicate that vaccination of hBLT mice with TBK1-trained DC induces specific and interconnected histological patterns of infiltrated CD8^+^ T cell responses in the spleen that are associated with the retention of HIV-1 infected cells in this organ, preventing the spread and progression of HIV-1 infection in peripheral organs from these mice.

**Figure 5.**
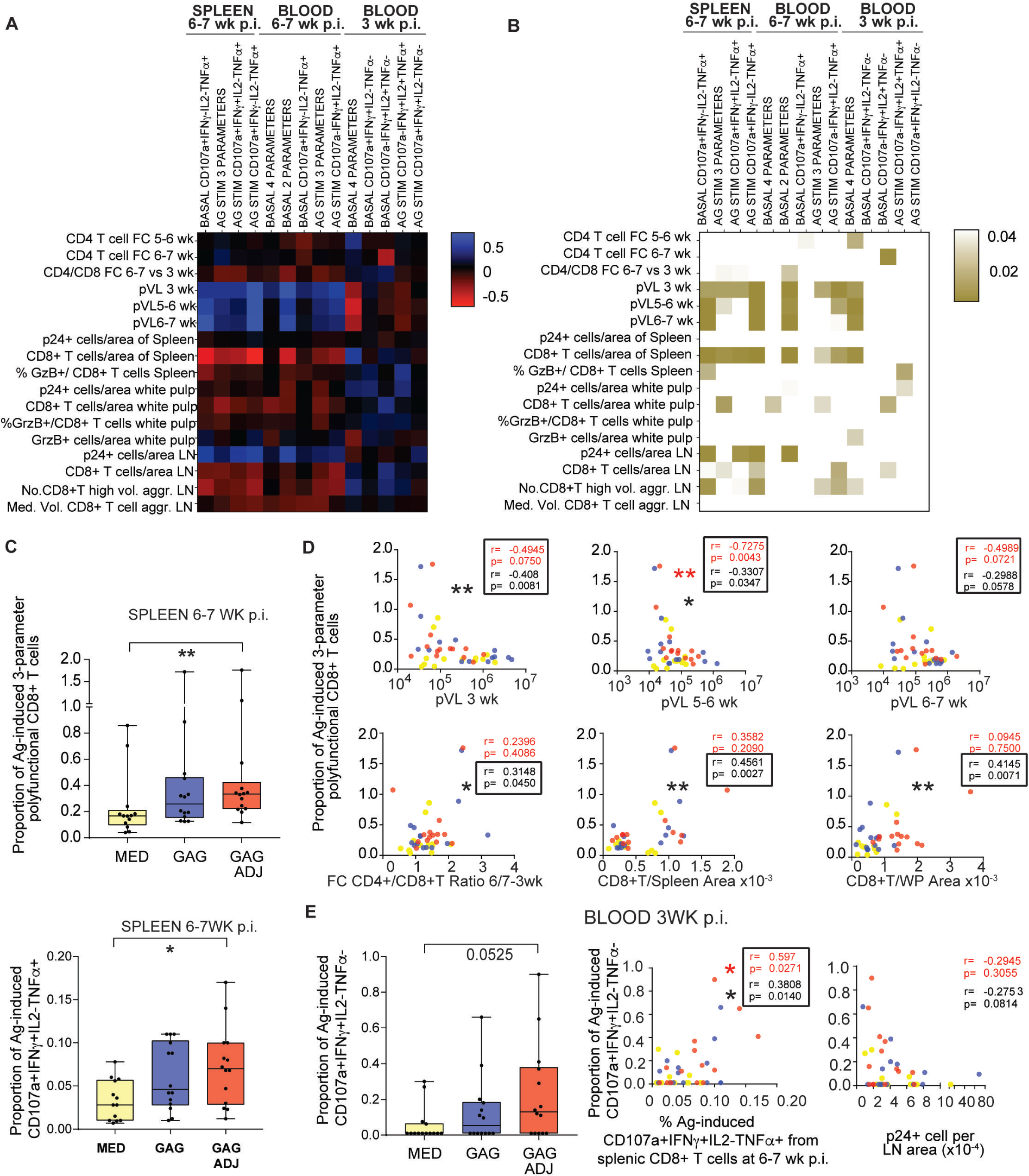
Identification of CD8^+^ T cell polyfunctional patterns associated with histological and clinical parameters of progression of HIV-1 infection in hBLT mice. (A-B): Correlation networks showing Spearman R (A) and p values (B) between selected clinical and histological parameters and basal and antigen-induced polyfunctional phenotype of splenic and circulating CD8^+^ T cell populations analyzed by flow cytometry at different times post-infection (B). Positive and negative correlations are highlighted in red and blue respectively; Significant p values for each correlation are highlighted in brown scale. (C): Proportion of splenic CD8^+^ T cells either co-expressing 3 out of 4 tested cytokine/degranulation parameters upon stimulation with a pool of HIV-1 Gag peptides (upper plot) or a splenic population of polyfunctional cells defined as CD107a^+^ INFγ^+^ TNFα^+^ IL-2^-^ detected under these conditions (lower plot). (D): Individual Spearman correlations between proportions of Gag-peptide induced splenic CD8^+^ T cells co-expressing 3 cytokine/degranulation parameters and the indicated clinical and histological parameters. Spearman p and r values are highlighted in each plot. *p<0.05; **p<0.01. (E): Proportions of CD107a^+^ INFγ^+^ TNFα^-^ IL-2^-^ included in circulating CD8^+^ T cells at 3weeks p.i. after HIV-1 Gag-peptide stimulation. Individual Spearman correlations between proportions of this population and Ag-induced 3 parameter polyfunctional cells and p24^+^ cell No. in LN are shown in the right. Significance for (C, E) was calculated using a Kruskal-Wallis multiple comparison test with Dunn’s correction (*p<0.05; **p<0.01).

### Polyfunctional CD8^+^ T cell responses in lymphoid tissue and blood from hBLT mice vaccinated with GAG-ADJ-DC

We next addressed the polyfunctional profiles of splenic and circulating CD8^+^ T cells from the three vaccinated hBLT mouse groups by analyzing expression of IFNγ, IL-2, TNFα and CD107a *ex vivo* and after stimulation with a pool of HIV-1 Gag peptides at 3 and 6/7 weeks post-infection. We analyzed the proportions of T cells co-expressing 2, 3 and 4 of the analyzed parameters as a readout for polyfunctionality and quantified all individual cell subsets by Boolean gating (Supplemental Figure 7). Overall, basal levels of cells displaying higher polyfunctionality tended to be increased on splenic and circulating CD8^+^ T cells from GAG-ADJ DC vaccinated mice, but differences did not reach statistical significance in all cases (Supplemental Figure 7A). In fact, only polyfunctional cells co-expressing two-parameters were significantly higher in the blood at 6/7 weeks p.i. of GAG-ADJ mice compared to GAG and MED control groups (Supplemental Figure 7C). In contrast, we observed a gradual increase in the induction of polyfunctional splenic CD8^+^ T cells co-expressing 3 and 2 out of 4 analyzed parameters after Gag peptide stimulation in the hBLT mice groups receiving GAG and GAG-ADJ DC vaccines. These differences were only significant in the GAG-ADJ hBLT group compared to MED mice (Supplemental Figure 7B; Figure 5C, upper plot). In fact, the increase of 3-parameter polyfunctional CD8^+^ T cells in GAG-ADJ hBLT mice after Gag peptide stimulation was driven by a more significant increase in the proportion of two specific subpopulations of CD107a^+^INFγ^+^IL2^-^TNFα^+^ and CD107a^-^ INFγ^+^ IL2^+^ TNFα^+^ CD8^+^ T cells, while the other combinations did not reach significance (Supplemental Figure 9A, Figure 5C bottom plot). To better determine whether the presence or the absence of total or specific polyfunctional populations was associated with reduced progression of HIV-1 infection, we performed a correlation network between these cell subsets and the clinical and histological parameters previously observed (Supplemental Figure 8). This unbiased approach identified histological and T cell responses significantly associated with viral control (Figure 5A-B, D; Supplemental Figure 8A-B). In the spleen, we did not observe any significant association of histological or clinical parameters with basal polyfunctional CD8^+^ T cell profiles differentially induced in the GAG-ADJ hBLT mice (Supplemental Figure 8A). However, significant associations between proportions of antigen-mediated induction of splenic polyfunctional CD8^+^ T cells co-expressing 3 parameters were found with less severe depletion of CD4^+^ T cell counts, infiltration of CD8^+^ T cells in the white pulp areas from the spleen and lower detection of infected p24^+^ cells in the LN (Figure 5D, Supplemental Figure 9B). Interestingly, when we analyzed which of the two Ag-induced 3-parameter polyfunctional T cells were more associated with virological, immunological and histological patterns, we observed that the population of CD107a^+^ IFNγ^+^ IL2^-^ TNFα^+^ CD8^+^ T cells significantly induced in GAG-ADJ hBLT mice was more significantly correlated with these parameters than the other CD107a^-^ INFγ^+^ IL2^+^ TNFα^+^ subset of CD8^+^ T cells induced in these animals (Figure 5C bottom panel; Supplemental Figure 9A-C). In particular frequencies of these CD107a^+^IFNγ^+^IL2^-^TNFα^+^ CD8^+^ T cells correlated more significantly with lower pVL at earlier time points (3 wk p.i; p=0.0081) and with higher CD4^+^/CD8^+^ T cell ratios in the blood (p=0.0468), higher infiltration of CD8^+^ T cells in the spleen (p=0.0010) and lower detection of p24^+^ cells in the LN (p=0.0030) at the time of sacrifice (6/7wk p.i) (Supplemental Figure 9A-C). These data suggest that CD8^+^ T cells from the spleen of GAG-ADJ hBLT mice display preserved abilities to induce specific patterns of polyfunctional cytotoxic and cytokine secreting cell subsets after antigen re-stimulation.

Our analyses also indicated that preserved polyfunctional responses after antigen stimulation from circulating cells were also associated with control of HIV-1 infection in hBLT mice. Interestingly, non-specific higher basal frequencies of polyfunctional CD107a^+^ IFNγ^+^ IL2^+^ TNFα^+^ in the absence of Ag stimulation and increased induction after Gag-stimulation of CD107a^-^ IFNγ^+^ IL2^+^ TNFα^+^ cells in circulating CD8^+^ T cells were not associated with protection parameters but seemed to be indicative of pronounced disease progression (Figure 5A-B, Supplemental Figure 9D). However, we found that the polyfunctional CD107a^+^IFNγ^+^IL2^-^TNFα^-^ cell population induced by Gag peptide stimulation from circulating CD8^+^ T cells at 3 weeks p.i. which was more significantly increased in GAG-ADJ hBLT mice (Figure 5E, left plot). Although proportions of these polyfunctional CD107a^+^IFNγ^+^IL2^-^TNFα^-^ at 3 weeks p.i. was not directly associated with clinical and histological parameters (Figure 5A-B), this subset significantly correlates with subsequent increased proportions of protective Ag-induced CD107a^+^IFNγ^+^IL2^-^TNFα^+^ (Figure 5E, right plots). Together, our results indicate that vaccination of hBLT mice with TBK1 trained DC enhance Ag-inducible precursors of polyfunctional T cell responses on circulating cells that can serve as biomarkers of tissue polyfunctionality and reduced progression of HIV-1 infection.

## Discussion

Our study evaluates the efficacy of DC simultaneously matured with two TBK1 adjuvants, a STING agonist and Poly I:C, inducing parameters of immune control of HIV-1 infection *in vivo*. We demonstrate that vaccination with TBK1-tailored DC is associated with reduced progression of HIV-1 disease in hBLT mouse model. Previous clinical trials evaluated the benefit of systemic administration of Poly I:C to HIV-1 infected individuals, and demonstrated an increase of HIV-1-specific T cell responses but the therapeutic benefit of this format remains unclear (41-43). In addition, while previous studies in a murine model suggested that HIV-1 vaccines administered systemically targeting DC via CD40 or DEC205 and Poly I:C as an adjuvant could induce antigen-specific responses (44-46), our work provides new proof of concept of beneficial effects of the administration of a TBK1-tailored DC vaccine in an *in vivo* humanized model without systemic adjuvant addition, which can trigger other cell populations. This is particularly relevant since it has been shown that systemic Poly I:C administration can lead to HIV-1 reactivation on CD4^+^ T cells (45, 47). Moreover, this study specifically explores the benefit of enhanced maturation of DC in the presence of a combination of Poly I:C and STING agonist, potentiating phosphorylation of TBK1 and IRF3 and more efficiently inducing the secretion of immunomodulatory cytokines such as IL-12 and IFNβ, which was associated with an increase of DC antigen presenting properties. However, we cannot completely rule out that in addition to activating TBK1, some of the adjuvants used in our study such Poly I:C could be also triggering additional pathways, which might also affect DC maturation. Despite this possibility, our data indicate that our combined adjuvant strategy is able to recapitulate some of the enhanced functional properties previously observed in DC from HIV-1 elite controllers (15, 48).

Importantly, while previous studies on HIV-1 vaccine prototypes have mainly focused on the phenotype or even polyfunctionality induced circulating T cells (49, 50), we were able to identify cellular and histological parameters associated with reduced spread of HIV-1 infection to secondary lymphoid organs, such as the spleen and the lymph nodes. Moreover, vaccination of hBLT mice with TBK1-tailored DC induced higher levels of infiltration of CD8^+^ T cells in white pulp areas of spleen, which were associated with accumulation of infected HIV-1 p24^+^ cells in these areas. This splenic phenotype was associated with higher volume of CD8^+^ T cell clusters and lower detection of infected cells in the lymph node of hBLT mice. These histological patterns bear some resemblance to follicular CD8^+^ T cell responses observed in primates able to control viral infection (26) and in HIV-1 controller patients (25). In fact, we observed expression of CXCR5 preferentially on CD8^+^ T cells infiltrating the white pulp areas from GAG-ADJ hBLT mice, which might support a follicular-like phenotype previously linked to viral control (51, 52). However, since deficiencies in lymphoid tissue architecture have been described in the hBLT model (32, 53), further characterization of white-pulp resident CD8^+^ T cells in the hBLT mouse needs to be conducted in order to better understand these potential similarities. In fact, in our study we did not address the causal relationships between the enrichment in cytotoxic CD8^+^ T cells in the red pulp and the infiltration of CXCR5^+^ CD8^+^ T cells in the white pulp and the differential accumulation of HIV-1 p24^+^ cells observed in these areas. Furthermore, the relationship between the observed histological distribution of splenic CD8^+^ T cells with inflammatory tissue fibrosis, previously linked to immunopathology of HIV-1 infection or the presence of CXCR5^+^ CD4^+^ T cells, was not addressed in our study and deserves further investigation (54, 55).

In addition to histological patterns, we identified in vaccinated hBLT mice preserved abilities of splenic CD8^+^ T cells to induce a polyfunctional population of tissue resident CD107a^+^ IFNγ^+^ TNFα^+^ IL2^-^ cells upon Ag stimulation that was associated with less severe depletion of circulating CD4^+^ T cells, higher infiltration of CD8^+^ T cells in the white pulp areas and lower numbers of infected p24^+^ cells in the lymph node, thus underscoring that these cells could display effective antiviral properties. Supporting this possibility, a number of studies have described that polyfunctional T cells co-expressing TNFα with other parameters correlate with protection against viral infections such as Zika and Cytomegalovirus (56, 57). However, future studies are needed to better understand the developmental kinetics and functional relationships of this particular subset of polyfunctional cells with other subpopulations that might also be present in the GAG-ADJ hBLT mice. While our data also indicate that TBK1-primed DC vaccination could induce control on plasma viral load, these effects could be mediated by HLA-variability or HIV-1 escape mutations induced in vaccinated hBTL mouse (33, 38). Although our study suggests that TBK-1 DC can induce multiple histological and immunological parameters associated with immune control of HIV-1 infection, we focused on analyzing them at key time points previously described to mark HIV-1 pathogenesis and detection of HIV-1 responses in the blood. Thus, further longitudinal studies with a larger number of hBLT mice and a broader range of time point analyses are required to better stablish the impact and evolution of the identified histological and immunological parameters during the course of HIV-1 infection and their relationship with protection.

Finally, an additional limitation of our study was the relatively high TCID50 dose and the administration route of HIV-1 to the hBLT mice studied. The primary objective of our study was to address whether vaccination of mice with TBK1-tailored DC could induce some level of protection against progression of HIV-1 infection in a model in which we ensured infection of all mice. However, new studies evaluating the beneficial effect of TBK1-tailored DC under more physiological conditions such as the use of lower viral titers and a mucosal administration route should be conducted. Despite these limitations, our study provides evidence of the beneficial effect of TBK1-tailored DC inducing more effective immune responses against HIV-1 at the histological, clinical, and cellular levels, and therefore it may be useful for the development of future vaccine strategies against HIV-1.

## Materials and Methods

### Isolation of human peripheral blood populations

Human Peripheral blood mononuclear cells (PBMC) were isolated by Ficoll (Pancoll, PAN Biotech) gradient centrifugation. Subsequently, conventional Dendritic Cells (cDC) and total T cells were purified from PBMC suspensions by negative immunomagnetic selection (purity >90%) using the Human Myeloid DC Enrichment Kit (STEMCELL) and the Untouched total human T cell (Invitrogen) kits, respectively. Monocyte-Derived Dendritic Cells (MDDC) were generated from adherent cells present in PBMCs and cultured in the presence of 100IU/ml of GM-CSF and IL-4 (Prepotech) for 5 days.

### In vitro functional assays

Human PBMC or purified primary cDCs were cultured in RPMI 1640 media supplemented with 10% Fetal Bovine Serum (HyClone) alone or in the presence of either 1μg/ml 2’3’-c’diAM(PS)2 (Invivogen) or 5μg/ml Poly I:C (SIGMA) or a combination of both for 24h. For all functional assays, stimulated cDC were washed with 1X PBS prior to the experiment. For mixed leukocyte reaction (MLR) assays, activated DC were co-cultured with allogenic T cells at a DC:T ratio of 1:2 in 96 round-bottom well plates for 5 days. At day 5, cultured lymphocytes were re-stimulated with 0.25μg/ml PMA (SIGMA) and ionomycin (SIGMA) for 1 h and cultured for 4 h in the presence of 0.5μg/ml Brefeldin A (SIGMA), 0.005mM Monensin and 0.2μg/ml anti-CD107a-APC antibody. Intracellular expression of INFγ and CD107a on cultured CD8^+^ and CD4^+^ T cells was then analyzed by flow cytometry. For the experiments evaluating *de novo*-priming of HIV-1 specific responses, total PBMCs from healthy donors were pre-stimulated with 5μg/ml of a pool of HIV-1 Gag peptides (NIH AIDS Reagent Program #11057) in the absence or the presence of the adjuvant combinations previously mentioned and kept in culture in media supplemented with 25 IU/ml IL-2 (Prepotech) for 2 weeks. Subsequently, cDC where isolated from PBMC from the same donor and activated under the same conditions in the absence or presence of apool of HIV-1 Gag peptides. After 16h, pre-cultured PBMC and stimulated autologous cDCs were co-cultured for additional 16h in the presence of Brefeldin A, Monensin and CD107a antibody and analyzed by flow cytometry as previously mentioned. All antibodies used for flow cytometry are listed in Table 1.

**Table 1.**
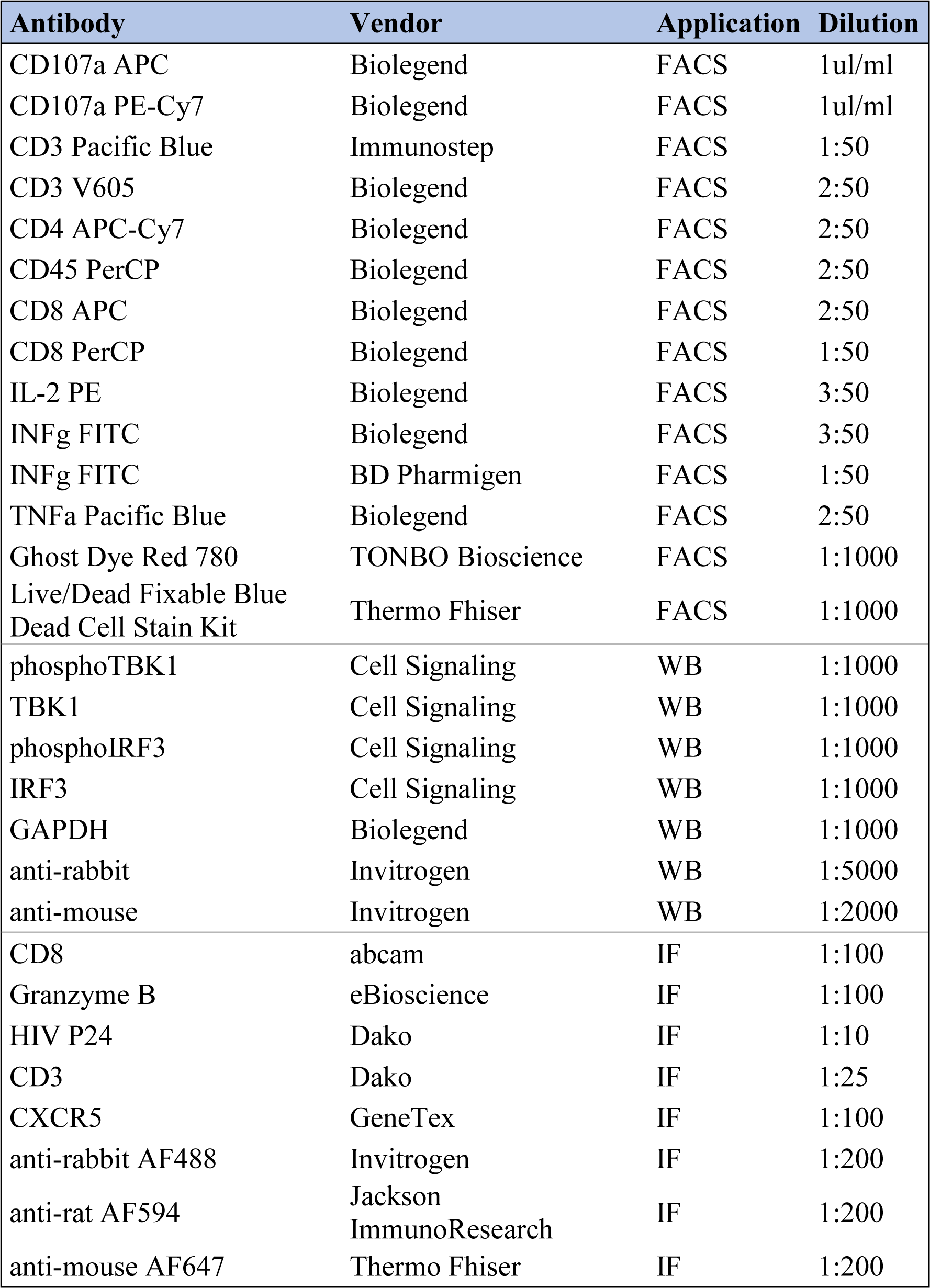
List of commercial antibodies used in the study

| **Antibody** | **Vendor** | **Application** | **Dilution** |
| --- | --- | --- | --- |
| CD107a APC | Biologend | FACS | 1ul/ml |
| CD107a PE-Cy7 | Biologend | FACS | 1ul/ml |
| CD3 Pacific Blue | Immunostep | FACS | 1:50 |
| CD3 V605 | Biologend | FACS | 2:50 |
| CD4 APC-Cy7 | Biologend | FACS | 2:50 |
| CD45 PerCP | Biologend | FACS | 2:50 |
| CD8 APC | Biologend | FACS | 2:50 |
| CD8 PerCP | Biologend | FACS | 1:50 |
| IL-2 PE | Biologend | FACS | 3:50 |
| INFg FITC | Biologend | FACS | 3:50 |
| INFg FITC | BD Pharmigen | FACS | 1:50 |
| TNFa Pacific Blue | Biologend | FACS | 2:50 |
| Ghost Dye Red 780 | TONBO Bioscience | FACS | 1:1000 |
| Live/Dead Fixable Blue<br>Dead Cell Stain Kit | Thermo Fhiser | FACS | 1:1000 |
| phosphoTBK1 | Cell Signaling | WB | 1:1000 |
| TBK1 | Cell Signaling | WB | 1:1000 |
| phosphoIRF3 | Cell Signaling | WB | 1:1000 |
| IRF3 | Cell Signaling | WB | 1:1000 |
| GAPDH | Biologend | WB | 1:1000 |
| anti-rabbit | Invitrogen | WB | 1:5000 |
| anti-mouse | Invitrogen | WB | 1:2000 |
| CD8 | abcam | IF | 1:100 |
| Granzyme B | eBioscience | IF | 1:100 |
| HIV P24 | Dako | IF | 1:10 |
| CD3 | Dako | IF | 1:25 |
| CXCR5 | GeneTex | IF | 1:100 |
| anti-rabbit AF488 | Invitrogen | IF | 1:200 |
| anti-rat AF594 | Jackson<br>ImmunoResearch | IF | 1:200 |
| anti-mouse AF647 | Thermo Fhiser | IF | 1:200 |

### Western blot analysis

Total protein lysates from MDDC and cDC cultured for 1 h in the presence of media or individual or combined Poly I:C and 2′3′-di AM(PS) agonists were obtained using RIPA buffer containing 1% phosphatase and protease inhibitors (Roche Diagnostics). Subsequently, protein lysates were resolved in a 10% agarose gel with SDS and transferred to a nitrocellulose membrane (Fisher Scientific). Membranes were blocked in 5% bovine serum albumin v (Sigma-Aldrich) in Tris buffered saline and incubated overnight with 1:100 dilution of primary anti-TBK1, anti-IRF3 or anti-GAPDH antibodies (Table 1). Then, membranes were incubated for 1 h with the appropriate anti-rabbit or anti-mouse secondary antibodies (Table 1). Protein band intensity was quantified by analyzing chemiluminescence detected using an ImageQuant 800 system (Amersham).

### Gene expression validation and RT-qPCR

RNA was isolated from cDC cultured in the absence or the presence of TBK1 adjuvants using RNeasy Micro Kit (Qiagen) according to manufacturer instructions. Subsequently, cDNA was synthesized using the Reverse Transcription kit (Promega). Transcriptional levels of IFN-β, IL-12 and TNFα were analyzed by semiquantitative PCR (SYBR Green assay Go Taq qPCR Master Mix; Promega) with specific primers (Metabion) on an Applied Biosystems StepOne Real-Time PCR system (Applied Biosystems). Relative gene expression was calculated after normalization to β-actin transcriptional levels.

### Generation of humanized BLT-mice

NOD/SCID IL2Ry-/- (NSG) mice transplanted with human bone marrow, fetal liver and thymus (BLT-mouse) were generated as previously described (32) at the Human Immune System Core from the Ragon Institute and Massachussets General Hospital in collaboration with Dr. Vladimir Vrbanac, Dr. Maud Deruaz and Dr. Alejandro Balazs. Mice were housed in microisolator cages and fed autoclaved food and water at a pathogen-free facility. Human immune reconstitution was monitored for 17 weeks and mice were considered reconstituted with proportions of human CD45^+^ lymphocytes superior to 30%.

### DC vaccination and HIV-1 infection of hBLT mice

Dendritic cells (DC) were generated from the same CD34^+^ HSC precursors used to reconstitute the corresponding batch of hBLT mice in the presence of 100IU/ml FLT3L, SCF, IL-7 and GM-CSF (Prepotech). After 10 days, cDC (CD14^+^ HLA-DR^-^) and MoDC (CD14^+^ HLA-DR^+^) present in cultures were sorted and incubated in media in the absence (MED mice group) or in the presence of 5ug/ml of a Gag pool of peptides (GAG mice group) alone or in combination with 1μg/ml of 2’3’-c’diAM(PS) and 5 μg/ml Poly I:C adjuvants (ADJ mice group). After 24 h, cDC and MoDC from each culture condition were pooled and hBLT mice were intravenously vaccinated in the tail vein with approximately 10^5^ total DC per animal. Two weeks after vaccination, hBLT mice were infected intravenously with a dose of 10,000 TCID50 of HIV-1JR-CSF. For histological analyses some unvaccinated uninfected mice were included as controls.

Plasma HIV-1 viral loads were assessed at 3 and 5/6 and 6/7 weeks post-infection by isolating viral RNA from plasma and quantified by RT-qPCR as previously described (38). Circulating CD4^+^ T cell counts were assessed at day 0, 3 weeks, 5/6 weeks and 6/7 weeks post-infection by flow cytometry using counting beads (CountBright, ThermoFisher).

### Histological analysis of tissue sections from hBLT mice

Lymph nodes and spleens were paraffin-embedded and segmented in fragments of 2 μm of thickness in a Leica microtome. Tissue sections deparaffinization, hydration and target retroviral were performed with a PT-LINK (Dako) previous to staining.

For paraffin-preserved tissue, we used rabbit anti-human CD8 (abcam), rabbit anti-human CXCR5 (GeneTex), rat anti-human CD8 (Bio-Rad), rat anti-human Granzyme B (eBioscience), mouse anti-human CD3 (Dako) and mouse anti-HIV-1 P24 (Dako), as primary antibodies; and goat anti-rabbit AF488 (Invitrogen), donkey anti-rat AF594 (Jackson ImmunoResearch) and donkey anti-mouse AF647 (ThermoFisher) as secondary antibodies. Images were taken with a Leica TCS SP5 confocal and processed with the LAS AF software. 3-D CD8^+^ T cell aggregations were analyzed with Imaris 9.1 software. CD8^+^ T cell, Granzyme B and HIV-1 P24 cell counts, co-localization and distance 2-Dimensions maps were analyzed with ImageJ software. In some cases, spleen tissue sections were also stained with hematoxylin and eosin to discriminate white (no eosin staining) and red pulp (intense eosin staining due to enrichment in erythrocytes) areas containing nucleated cells (hematoxylin stained).

### Analysis of polyfunctional T cell responses

Blood was extracted from hBLT mice at 3 and 5-6 weeks post-infection and lysed with Red Blood Cell Lysis Buffer (SIGMA). T cells were activated for 1.5 h with 5μl/ml of anti-CD28 and anti-CD49d in the presence or absence of 6.4μg/ml of a Gag pool of peptides in the presence of 0.5μg/ml Brefeldin A, Golgi Plug and CD107a antibody (see Table 1). After 5 h of incubation, polyfunctionality of T cell response was assessed by INFγ, IL2, TNFα and CD107a expression by multicolor flow cytometry panel (all antibodies used are listed in Table 1) in a BD LSR Fortessa Instrument (BD Biosciences). Polyfunctionality was evaluated using Boolean gating obtained with FlowJo v10 software.

### Statistics

Significance of phenotypical and functional differences between paired conditions or different animals were assessed using a Wilcoxon matched-pairs signed-rank test or Mann-Whitney U test, or using a Kruskal-wallis or Friedman test followed by a Dunńs post-hoc multiple comparison test, as appropriate. Dependence of contingency tables values were calculated with Chi-square statistic. Association between clinical, histological and phenotypical parameters were calculated using non-parametric Spearman correlation individually between two parameters or using a correlation network. All statistical analyses were performed using the GrapPad Prism 8 Sofware.

### Ethics statement

## Supporting information

Combined Supplemental Figures

## Funding

EMG was supported by the NIH R21 program (R21AI140930), the Ramón y Cajal Program (RYC2018-024374-I), the MINECO/FEDER RETOS program (RTI2018-097485-A-I00) and by Comunidad de Madrid Talento Program (2017-T1/BMD-5396). MJ.B is supported by the Miguel Servet program funded by the Spanish Health Institute Carlos III (CP17/00179), the MINECO/FEDER RETOS program (RTI2018-101082-B-100) and Fundació La Marató TV3 (201805-10FMTV3). EMG and MJB are funded by “La Caixa Banking Foundation H20-00218). M.C.M was also funded by the NIH (R21AI140930). FS-M was supported by SAF2017-82886-R from the Ministerio de Economía y Competitividad and HR17-00016 grant from “La Caixa Banking Foundation. We also would like to thank the NIH AIDS Reagent Program, Division of AIDS, NIAID, NIH for providing HIV-1 PTE Gag Peptide Pool from NIAID, DAIDS (cat #11057) for the study. Finally, the Microscopy Unit from Centro Nacional de Investigaciones Cardiovasculares provided technical support for the microscopy image processing and analysis.

## Author contributions

E.M.G., V.V., D.C., M.D., A.B., M.J.B. developed the research idea and study concept, designed the study and wrote the manuscript; E.M.G., V.V. supervised the study; M.C.M., D.C. M.D and S.T. designed and conducted most experiments and equally contributed to the study; T.A. and D.C. provided longitudinal VL data evolution in BLT mice from a pilot experiment M.C.M. performed the histology staining and the image analysis of tissue sections from the study. M.D and D.C. provided critical feedback during experimental design and execution phases of the studies and were directly involved in the experiments.

M.J.B. and C.S. provided reagents and support for the histological analyses performed in the study. F.S.M., A.A., M.A.MF, I.D.S, L.G.F and J.S. provided peripheral blood, reagents and participated on the analysis of the data.

## Declarations of Interests

The authors declare no competing interests.

## Supplemental figure legends

**Supplemental figure 1. Impact of TBK1 adjuvants in activation and function of cDC *in vitro*.** (A): Representative Western blot analysis of TBK1 and IRF3 phosphorylation in primary cDCs cultured for 1 h in the presence of media alone or with 2’3’-c-di-AM(PS)2 and/or Poly I:C. Ratios for phosphorylated vs total TBK1 and IRF3 proteins are shown on the right. Significance was calculated using a Kruskal-Wallis multiple comparison test with Dunn’s correction (*p<0.05). (B): Flow cytometry analysis of Mean Fluorescence Intensity (MFI) of CD40 (left) and CD86 (right) in cDC following culture in the absence or the presence of different indicated adjuvant combinations (n=8 experiments). Significance was calculated using a two-tailed Wilcoxon test (*p<0.05). (C): Flow cytometry dot plots showing analysis of IFNγ versus CD8 on gated CD8^+^ T cells from healthy individuals exposed to autologous cDCs pre-cultured in media alone or activated with 2’3’-c-di-AM(PS)2 and Poly I:C in the absence or the presence of a pool of HIV-1 Gag peptides. Dot plots from three representative experiments are shown. Number below gates represent proportion of positive cells.

**Supplemental figure 2. *In vitro* generation and isolation of HSC-derived DC and experimental design for *in vivo* hBLT vaccination and analysis.** (A): Representative pre-sorting gaiting strategy showing cell populations derived from human fetal CD34^+^ HSC cultured *in vitro* for 2 weeks (see methods). Conventional dendritic cells (cDC) and monocyte derived DC-like (MoDC-like) derived from HSC were defined as live CD33^+^ HLA-DR^+^ myeloid cells differing on CD14 expression, respectively. (B): Flow cytometry analysis of CD14 vs HLA-DR expression on sorted cDC (upper plots) and MoDC-like cells (lower plots). Proportion of cells included on each gate are highlighted. Levels of CD11c expression overlayed with FMO controls (blue histograms) for each of these two populations (red histograms) is also shown on the right. (C): Flow cytometry analysis of expression of CD40 versus CD86 on sorted CD34^+^ HSC-derived cDC and MoDC-like cultured in just media (MED) or in the presence of a pool of HIV-1 Gag peptides alone (GAG) or in combination with the TBK1 adjuvant cocktail (GAG-ADJ). Numbers in quadrants indicate proportions of positive cells. (D): Schematic representation of the experimental generation of hBLT mice, *in vivo* vaccination regime and analysis design. (E): Analysis of hBLT mice weight during the course of the experiment. Individual weights of hBLT mice are shown in a lighter color and median for each hBLT mouse subgroup is highlighted in a darker color and thicker lines (yellow for Media (MED), blue for Gag pool (GAG) and red for Gag pool + adjuvants (GAG AGJ)).

**Supplemental figure 3. Evolution of plasma HIV-1 viral loads in vaccinated hBLT mice infected with HIV-1.** (A): Pilot experiment showing RT-qPCR analysis of the evolution of plasma HIV-1 (RNA copies/ml) in n= 7 hBLT mice at different weeks after infection with JRCSF HIV-1 virus. (B-C): RT-qPCR analysis of plasma viral load in hBLT-mice vaccinated with MED, GAG and GAG-ADJ treated DCs at 3, 5/6 and 6/7 weeks post infection (B) and stratified in CD4 Hi and CD4 Low animals included within each hBLT mouse subgroup at 6/7 weeks post infection (C). Pie charts (lower panel C) represent the proportions of mice presenting plasma viral load either equal or higher than 10^5^ copies/ml (dark color) or lower than 10^5^ copies/ml (light color). Statistical significance of differences was calculated using a Chi-square test with Yates correction (****p<0.0001).

**Supplemental figure 4. Depletion of circulating CD4+ T cells in vaccinated hBLT mice infected with HIV-1.** (A-C): Fold-change in peripheral hCD4^+^ T cell counts at 3, 5-6 and 6-7 weeks post infection with HIV-1 in the study (A) or shown individually in two separate experiments performed with different batches of hBLT mice (C; experiment 1, n=24, left and experiment 2, n=18, right). Individual data for each mouse was normalized to the corresponding baseline hCD4^+^ T count values present at day 0. (B): Fold change in CD4^+^/CD8^+^ T cell ratios in the blood at 5/6 and 6/7 weeks post-infection from the values observed at 3 weeks post-infection in the indicated groups of vaccinated animals. Statistical significance was calculated using a two-tailed matched-pairs Wilcoxon test (*p<0.05; **p<0.01).

**Supplemental figure 5. Histological analysis of hBLT mice splenic architecture and association with CD8^+^ T cell activation.** (A): Representative image of a hematoxylin-eosin staining of a full spleen section from a hBLT mouse used for the study and defining white and red pulp areas (magnification 5x). Dashed areas include white pulp and exclude red pulp and (i) and (ii) sections from these regions are further zoomed in the lower panels (magnification 20x). (B): Quantification of percentage of cytotoxic Granzyme B^+^ hCD8^+^ T cell from total splenic hCD8^+^ T cells (upper panel) and number of HIV p24^+^-infected cells per square microns (lower panel) detected per splenic section of the hBLT mice from the indicated subgroups. Significance was calculated using a Kruskal-Wallis multiple comparison test with Dunn’s correction. (C): Analysis of the percentages of cytotoxic Granzyme B^+^ cells from total CD8^+^ T cells found in the white pulp (WP, upper plot) and in the red pulp (RP, lower plot) areas in the spleen of the indicated hBLT mouse subgroups. Significance between white and red pulp paired values was calculated using a two-tailed Wilcoxon test (*p<0.05; **p<0.01; ***p<0.001). Intergroup significance was calculated using a Kruskal-Wallis multiple comparison test with Dunn’s correction (*p<0.05). (D): Spearman correlation between proportions of Granzyme B^+^ CD8^+^ T cells present in the red versus the white pulp of the spleen of vaccinated hBLT mice. Spearman R and p values are highlighted in the upper right area of the plot. (E): Frequencies of CD8^+^ T cells and p24^+^ cells per spleen area in vaccinated MED (yellow), GAG (blue) and GAG-ADJ (red) hBLT mice stratified by less severe (CD4Hi) and marked (CD4Lo) depletion of circulating CD4^+^ T cell counts at 5/6 wk p.i. Statistical significance between values from CD4Hi ADJ-GAG mice and the indicated subgroups were performed using a two-tailed Mann Whitney test. (F): Representative confocal microscopy images (magnification 40x) from white pulp areas of the spleen of a GAG (left panel) and a GAG-ADJ (right panel) spleen section stained with anti-CD3 (yellow), anti-CD8 (green), CXCR5 (magenta); Nuclei were stained with DAPI (blue). White arrows highlight CXCR5^+^ CD8^+^ T cells in the white pulp areas from the spleen.

**Supplemental figure 6. Analysis of CD8^+^ T cell clusters in the lymph node from hBLT mice.** (A): Analysis of the volume of CD8^+^ T cell clusters detected on hBLT LN tissue sections using the Imaris 9.2 software. CD8^+^ clusters are colored with a gradient from higher volumes (red and orange) to lower volumes (purple and dark blue). (B): Proportion of infected HIV-1 p24^+^ cells per lymph node area stratified by CD4 Hi and CD4 Low animals included on each hBLT mouse subgroup. (C): Individual Spearman correlations of p24^+^ cells per LN area versus plasma viral loads at different time points and CD8^+^ T cell per spleen area at 6/7 weeks p.i. Values of r and p in Total (black) and GAG-ADJ (red) hBLT mice are highlighted on each plot. *p<0.05; **p<0.01; ***p<0.001; ****p<0.0001. (D): Quantification of numbers of large (500-3000000 μm^3^, upper pot; red line showing high-volume elements cut-off) and low (0-500 μm^3^, lower pot) volume CD8^+^ T cell clusters obtained with the Imaris 9.2 software for every single lymph node and per hBLT mouse.

**Supplemental figure 7. Quantification of basal and HIV-1 peptide induced polyfunctional profiles of splenic and circulating CD8^+^ T cells from hBLT mice.** (A, B): Percentages of polyfunctional splenic CD8^+^ T cells at 6/7 weeks post-infection defined as lymphocytes co-expressing either 4, 3 or 2 analyzed cytokine and degranulation parameters on gated CD8^+^ T cells either basally (A) or upon HIV-1 Gag peptide stimulation (B). (C-F): Percentage of polyfunctional cells, as previously defined, included on circulating CD8^+^ T cells at 3 and 6/7 weeks post-infection either basally (C for 6/7 wk p.i., and E for 3 wk p.i.) and after HIV-1 Gag peptide-stimulation (D for 6/7 wk p.i., and F for 3 wk p.i.). Statistical significance was calculated using a Kruskal-Wallis multiple comparison test with Dunn’s correction (*p<0.05; **p<0.01; ***p<0.001).

**Supplemental figure 8. Association of HIV-1 Gag-peptide induced splenic CD107a^+^ INFγ^+^ TNFα^+^ CD8^+^ T cells and HIV-1 disease progression parameters.** (A-B): Correlation network showing Spearman R (A) and p values (B) between the indicated clinical, histological, and basal and antigen-induced polyfunctional phenotype of splenic and circulating CD8^+^ T cell populations analyzed by flow cytometry at different times post-infection. Positive and negative correlations are highlighted in red and blue respectively; Significant p values for each correlation are highlighted in brown scale

**Supplemental Figure 9. Association of basal and HIV-1 Gag-peptide induced polyfunctional CD8^+^ T cells populations and histological and HIV-1 disease progression parameters. A):** Proportions of CD107a^+^IFNγ^+^IL2^+^TNFα^-^, CD107a^+^IFNγ^-^IL2^+^TNFα^+^ and CD107a^-^ IFNγ^+^IL2^+^TNFα^+^ 3-parameter polyfunctional subpopulations from splenic CD8+ T cells induced after HIV-1 Gag peptide stimulation CD8^+^ T cells. Statistical significance was calculated using a two-tailed Mann Whitney test, **p<0.01 (B-D): Individual Spearman correlation between proportions of CD107a^+^ INFγ^+^ IL2^-^ TNFα^+^ (Upper rows) and CD107a^-^ INFγ^+^ IL2^+^ TNFα^+^ (bottom rows) from splenic CD8^+^ T cell detected after HIV-1 Gag-peptide stimulation and the indicated virological (B) and immunological (C) parameters. Correlations between indicated histological and immunological parameters and proportions of CD107a^+^IFNγ^+^IL2^+^TNFα^+^ cells basally present in circulating CD8^+^ T cells at 3 weeks post-infection are shown in panel D. Spearman R and p values of all and ADJ-GAG hBLT mice groups are highlighted in black and red, respectively.

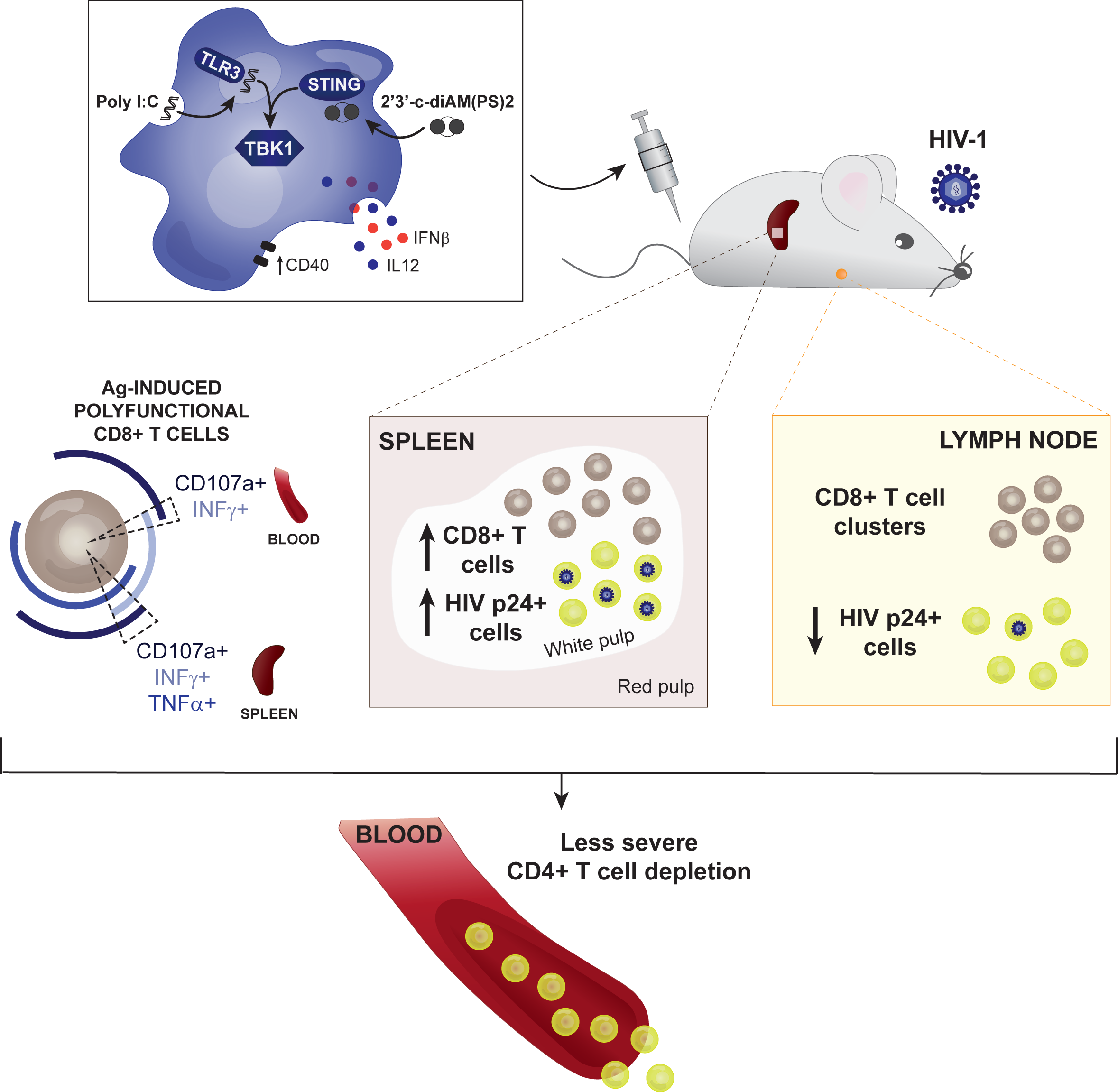

## Notes

### Competing Interest Statement

The authors have declared no competing interest.

