## Supplementary material for "TAILORED DC INDUCE PROTECTIVE HIV-1 SPECIFIC POLYFUNCTIONAL CD8+ T CELLS IN THE LYMPHOID TISSUE FROM HUMANIZED BLT MICE": Combined Supplemental Figures

### Supplementary figure 1

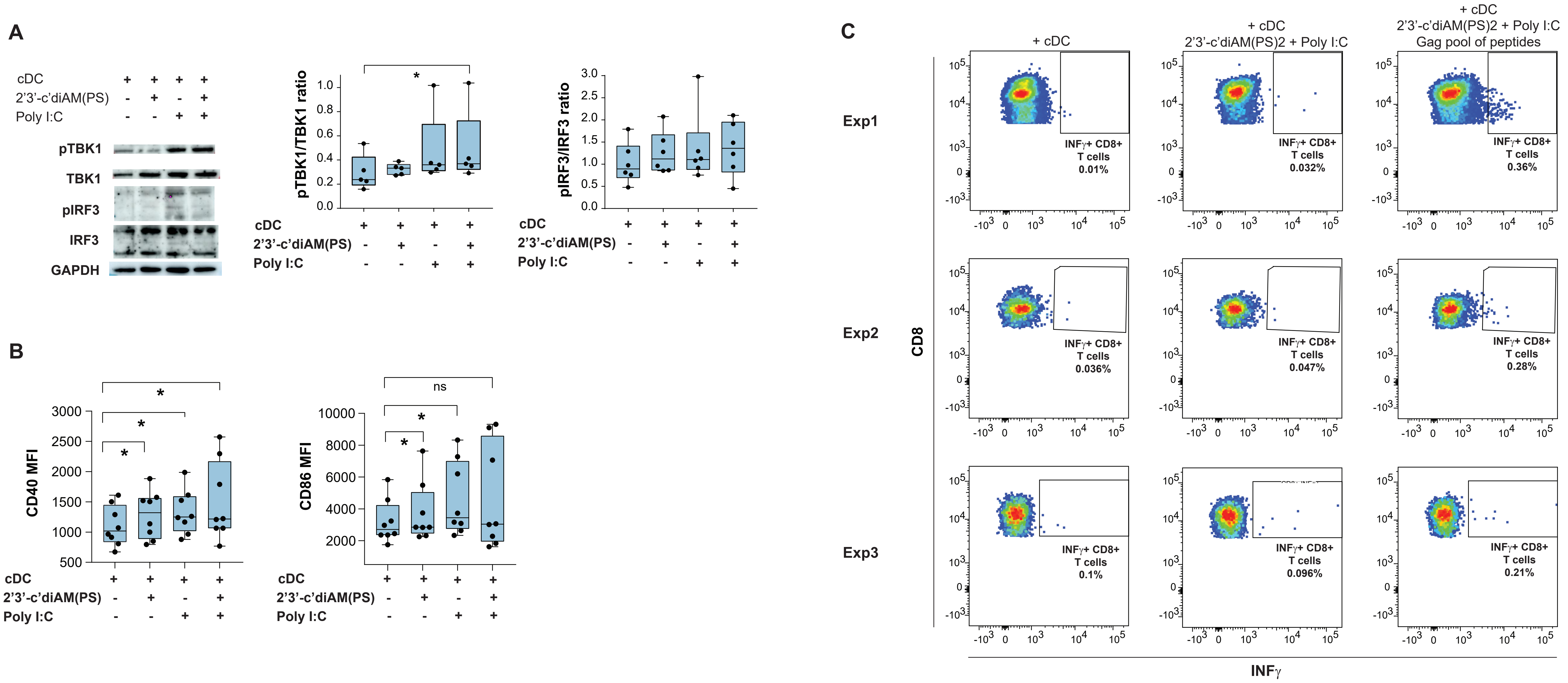

### Supplementary figure 2

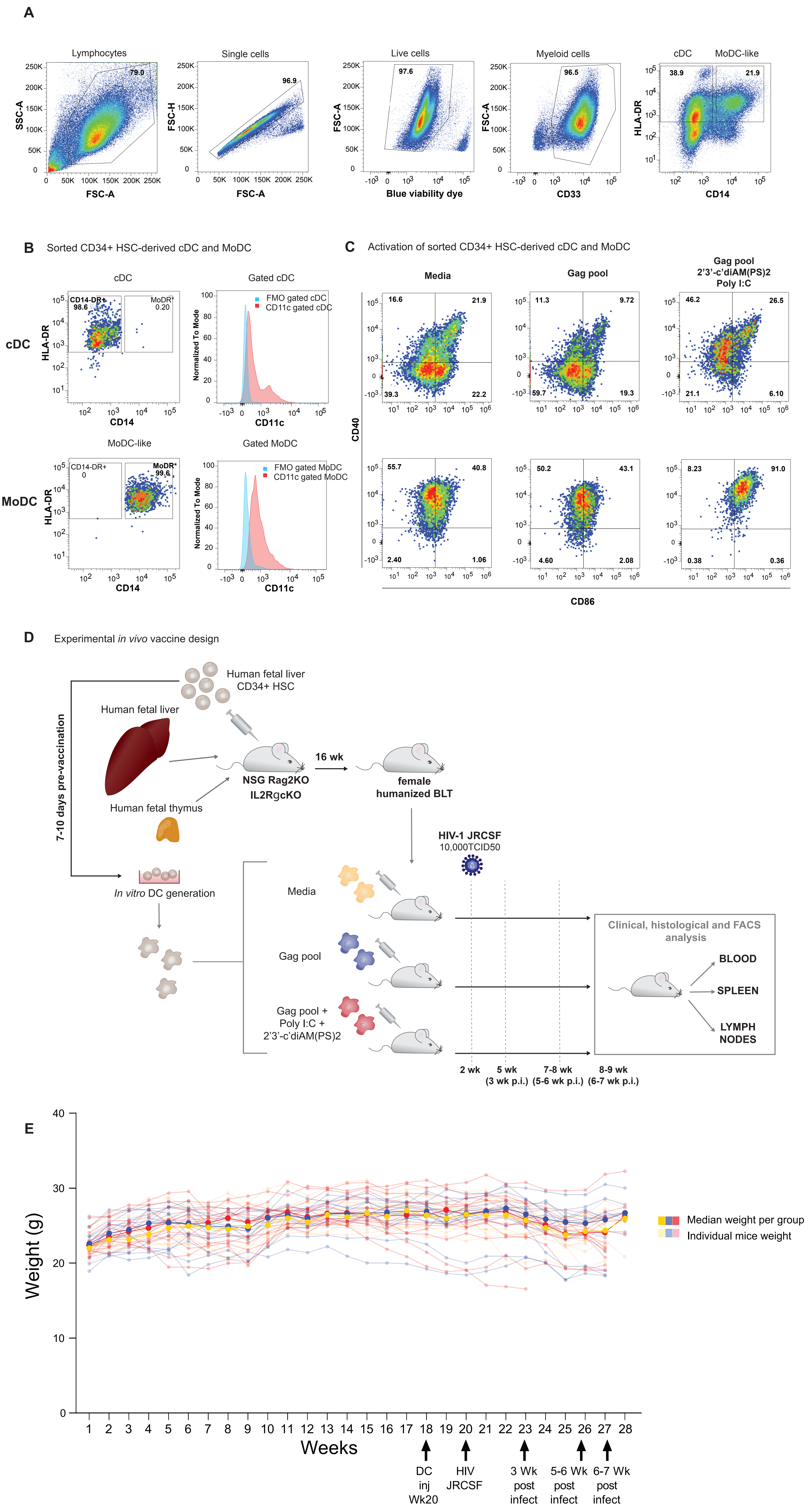

**Supplementary figure 3**

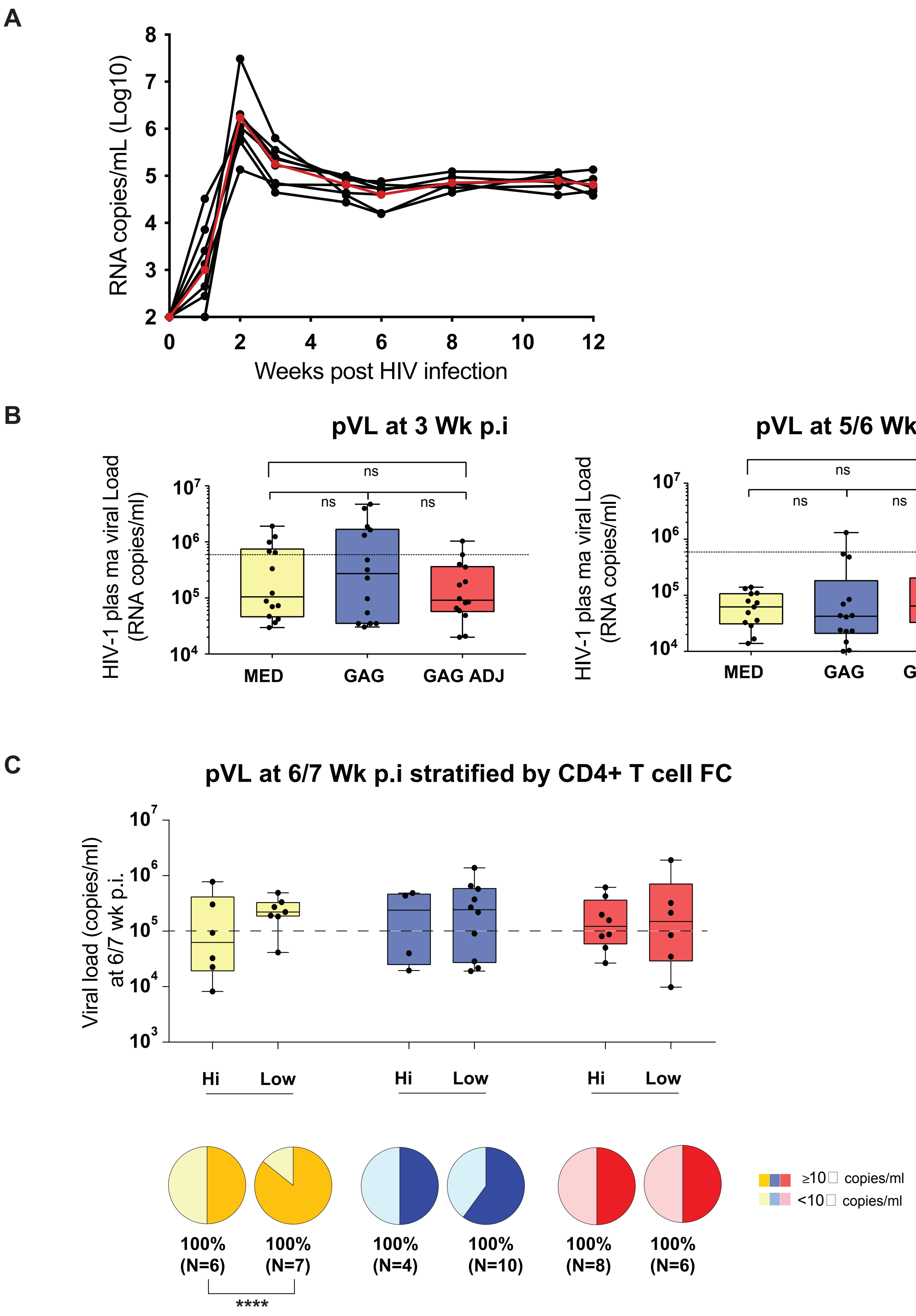

### Supplementary figure 4

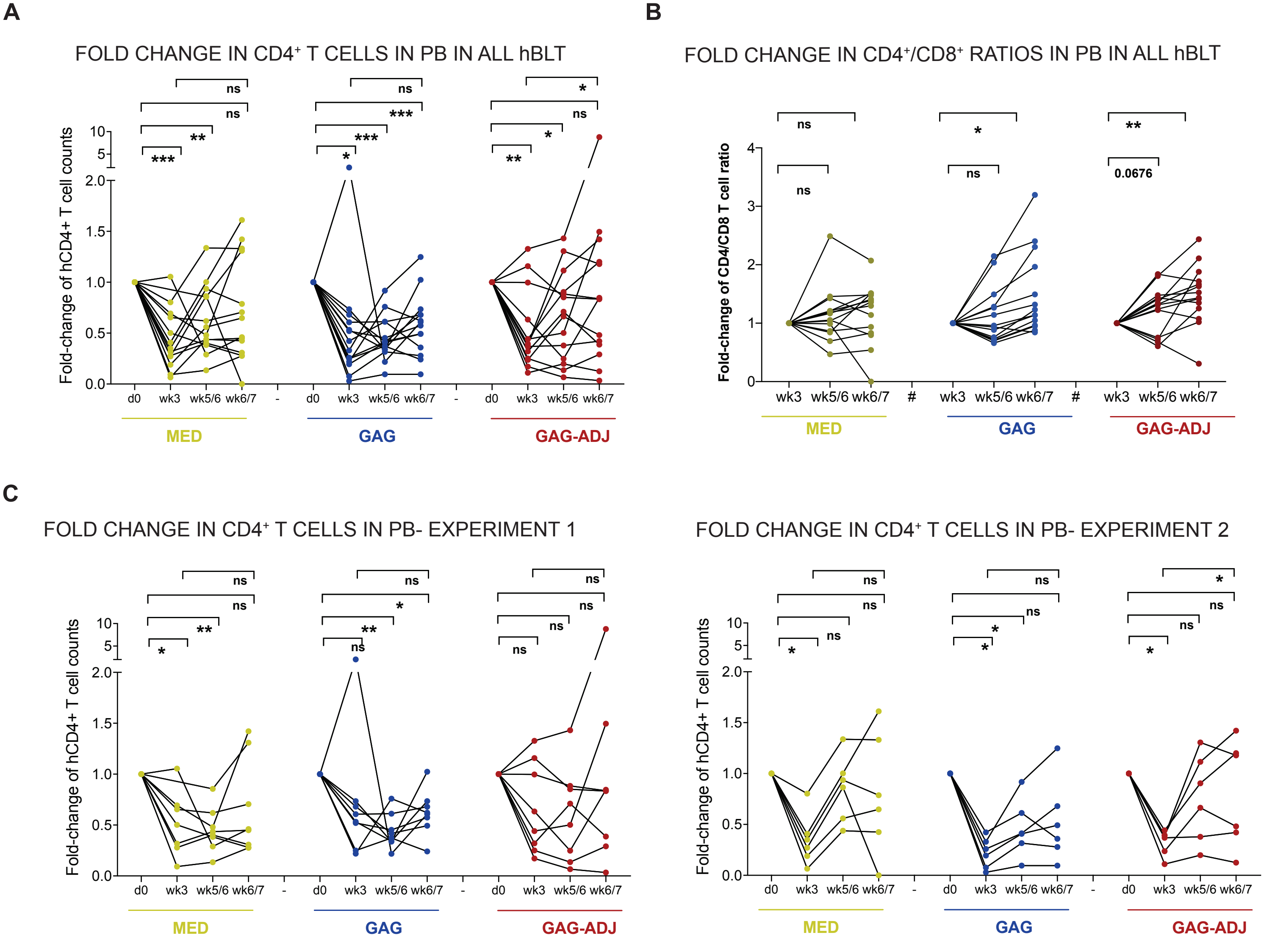

#### Supplementary figure 5

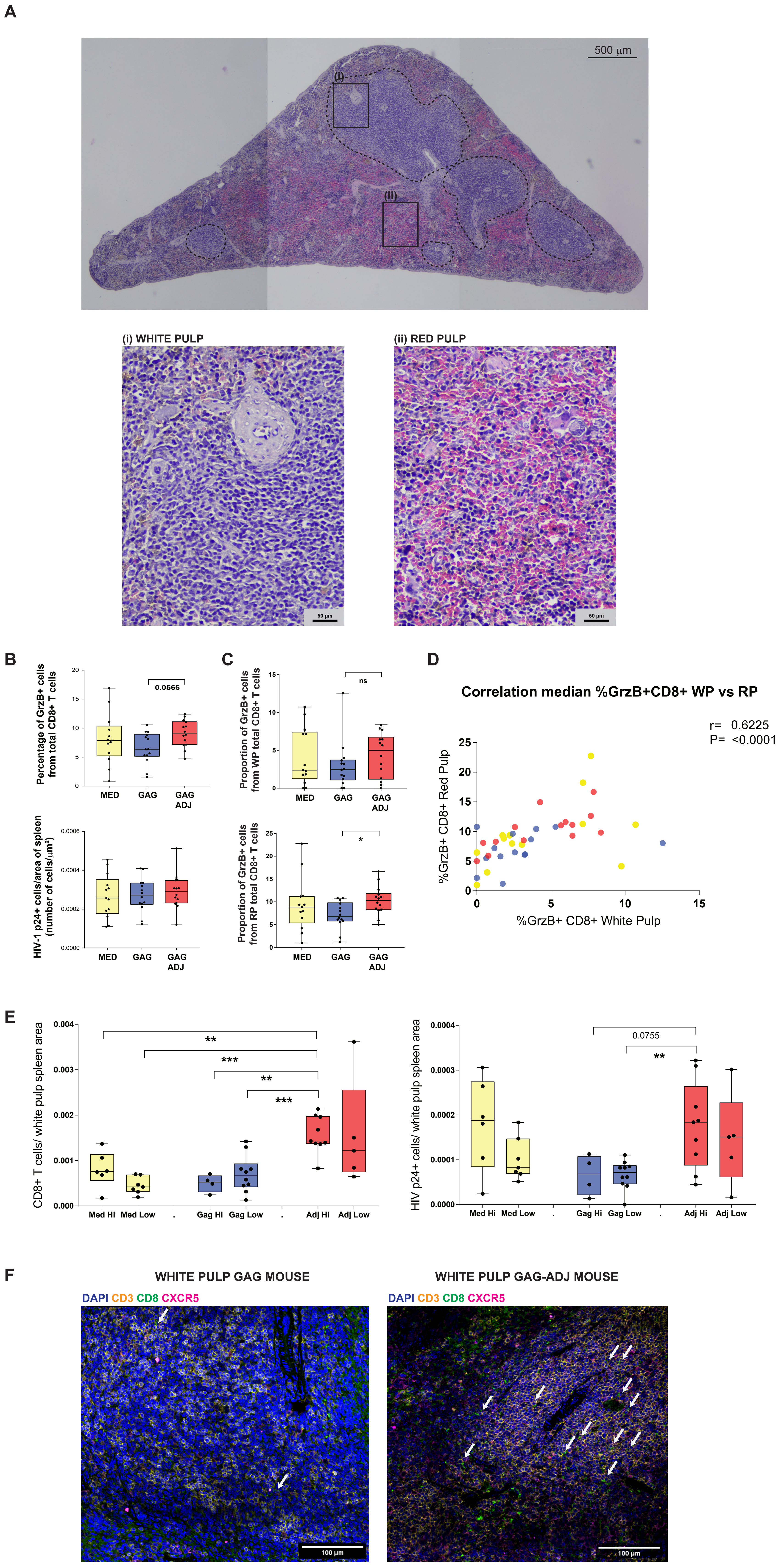

### Supplementary figure 6

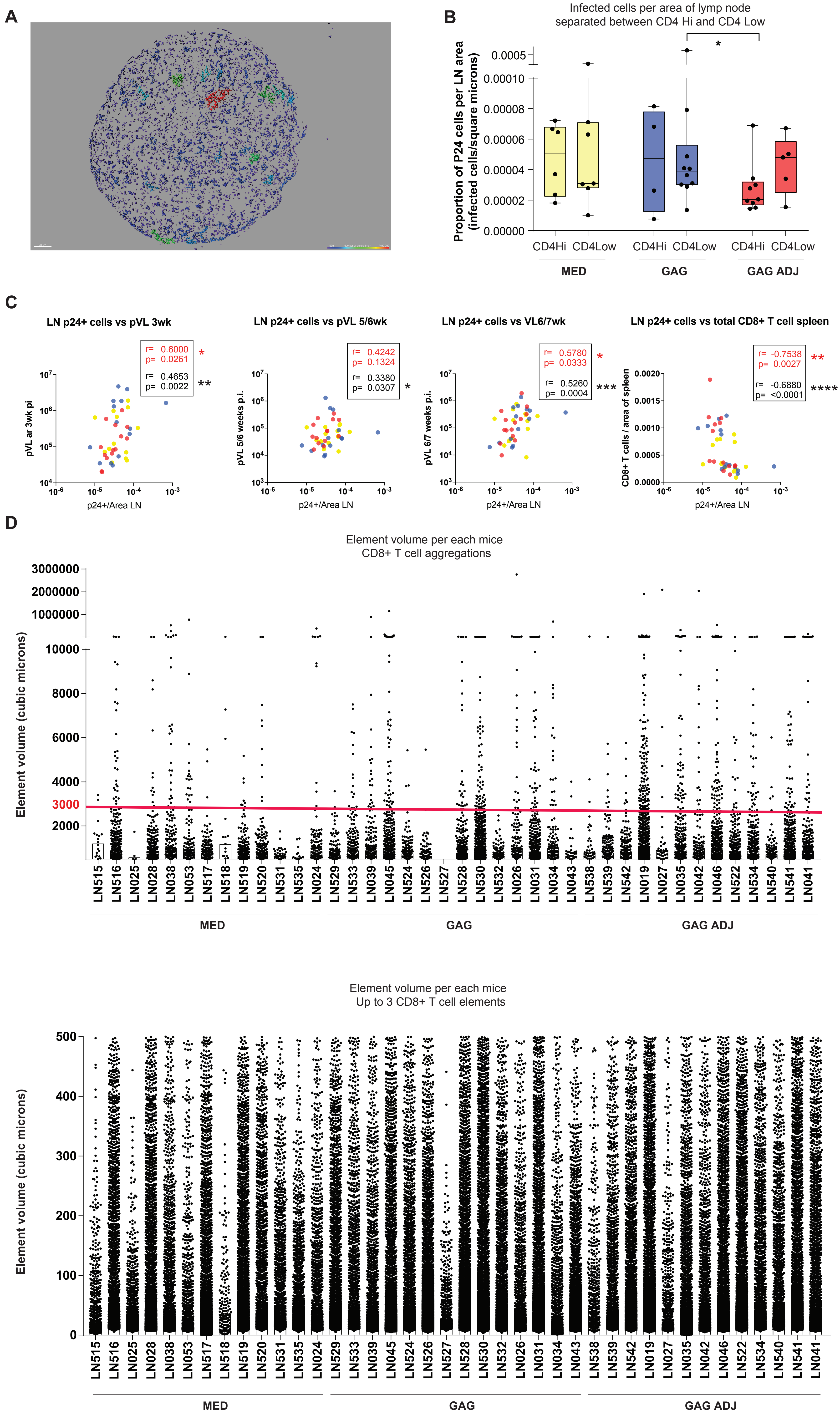

### Supplementary figure 7

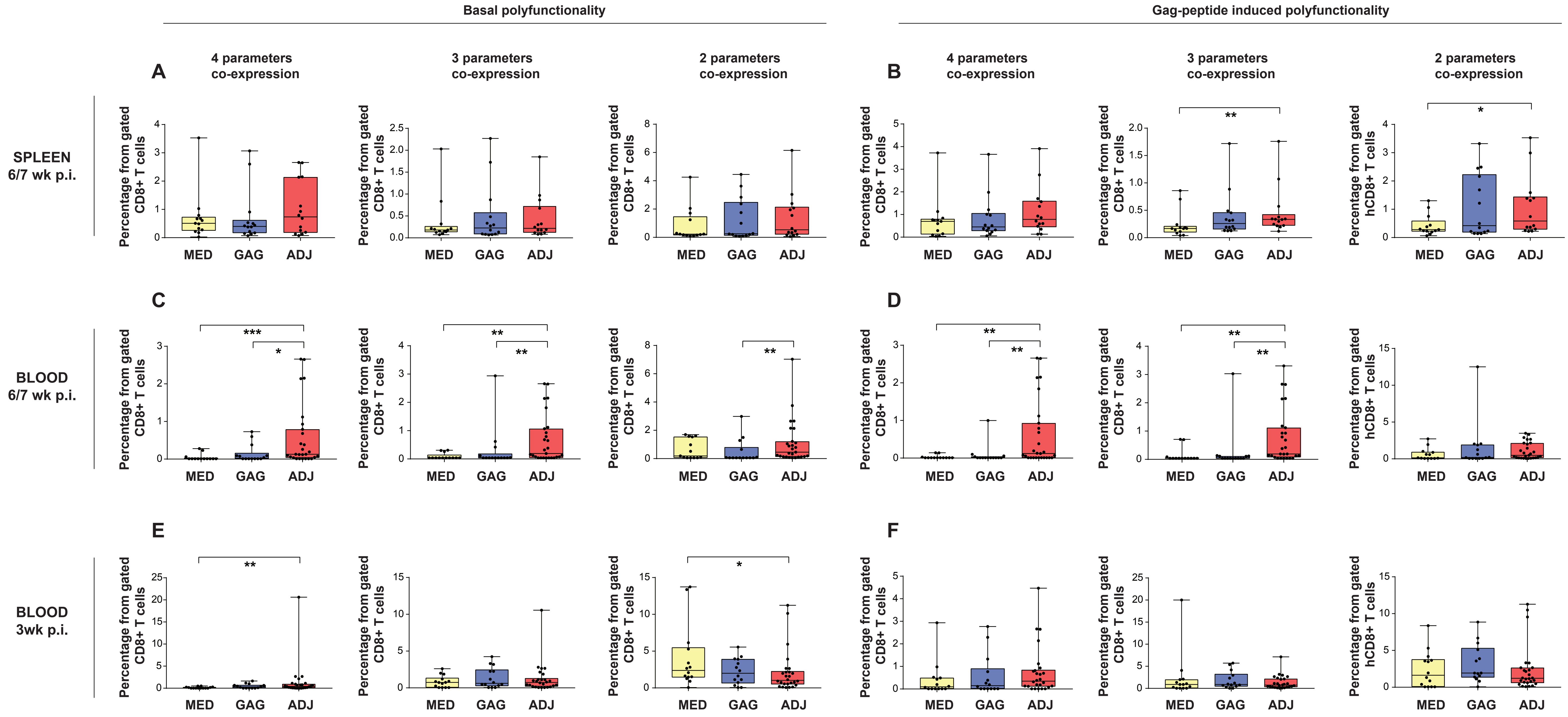

### Supplementary figure 8

A

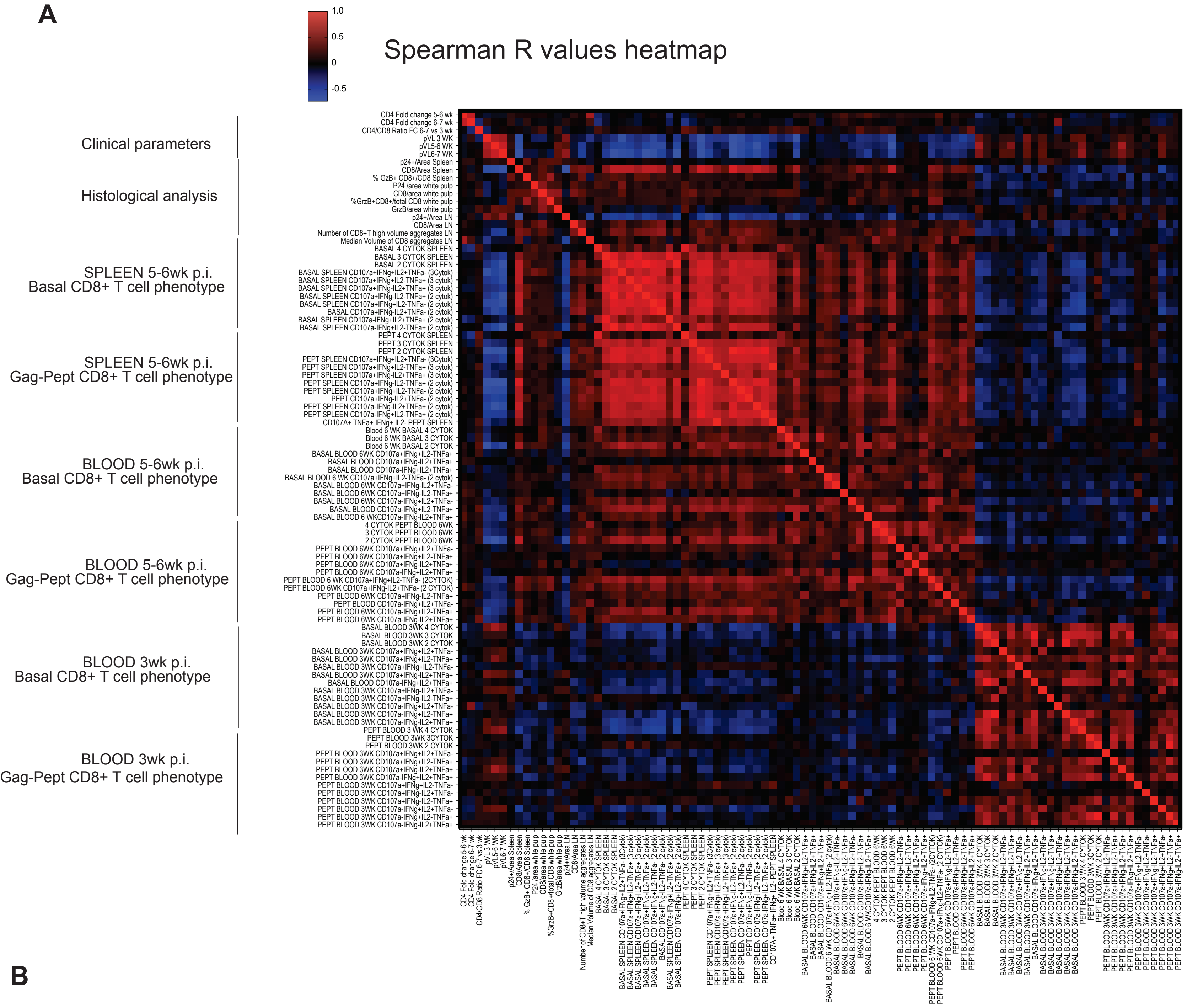


B

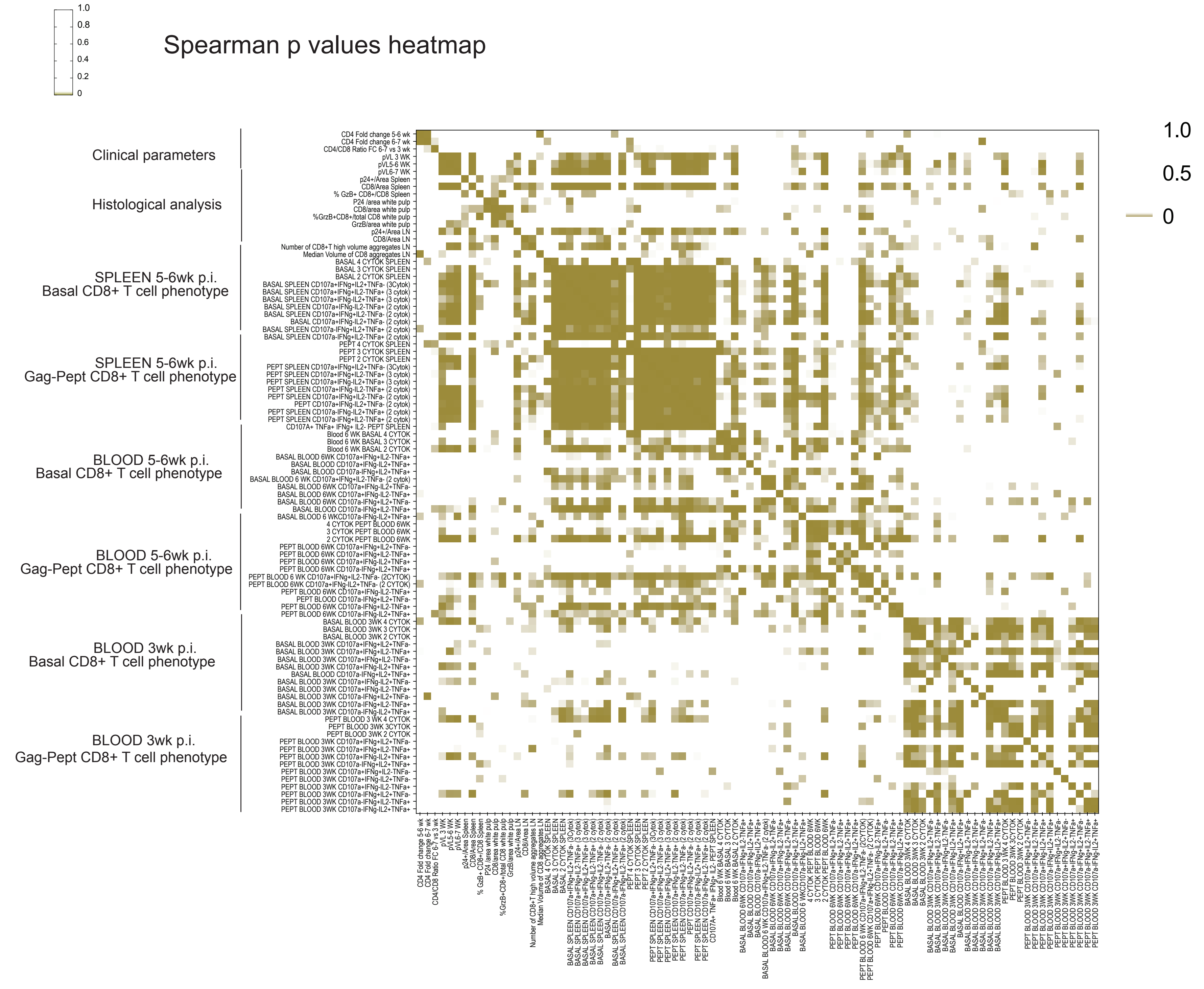

#### Supplementary figure 9

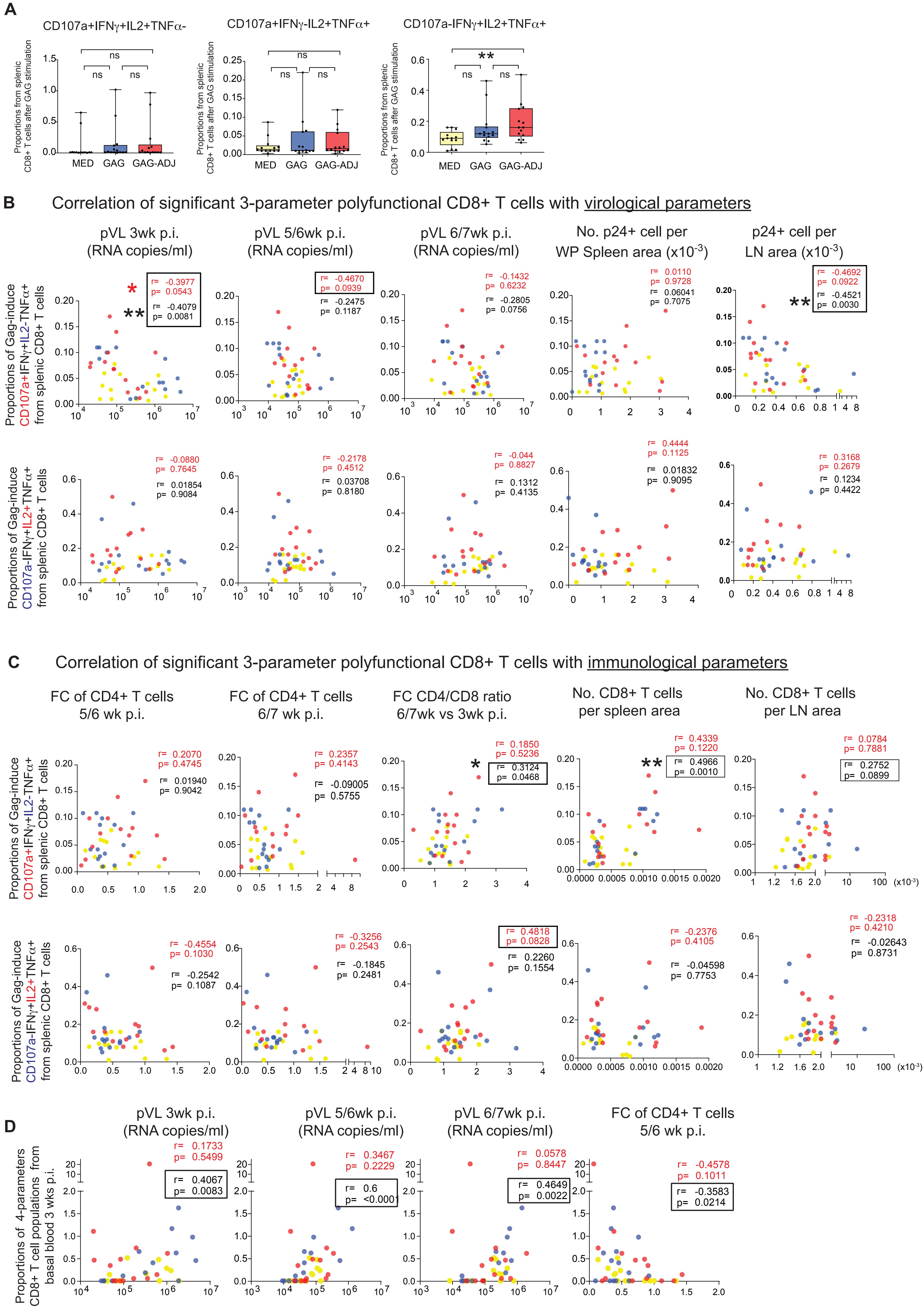
